# Polygenic predictions of occupational status GWAS elucidate genetic and environmental interplay for intergenerational status transmission, careers, and health

**DOI:** 10.1101/2023.03.31.534944

**Authors:** Evelina T. Akimova, Tobias Wolfram, Xuejie Ding, Felix C. Tropf, Melinda C. Mills

## Abstract

Socioeconomic status (SES) impacts health and the life course. This GWAS on sociologically informed occupational status measures (ISEI, SIOPS, and CAMSIS) using the UKBiobank (N=273,157) identified 106 genetic variants of which 8 are novel to the study of SES. Genetic correlation analyses point to a common genetic factor for SES. Within-family prediction and its reduction was attributable in equal parts to genetic nurture and assortative mating. Using polygenic scores from population predictions of 5-8%, we, firstly, showed that cognitive and non-cognitive traits – including scholastic and occupational motivation and aspiration – link genetic scores to occupational status. Second, 62% of the intergenerational transmission of occupational status can be ascribed to non-genetic inheritance (e.g., family environment). Third, the link between genetics, occupation, and health are interrelated with parental occupational status confounding the genetic prediction of general health. Finally, across careers, genetic prediction compresses during mid-career with divergence in status at later stages.

## Main

Socio-economic status (SES) is a core factor stratifying society, with deep impacts on wealth,^1^ health,^2^ family, and life course.^3^ Various disciplines, including economics, demography, public health, and sociology, have studied and operationalized this multidimensional construct, focusing on the ‘big three’ indicators: educational attainment, income and earnings, and occupational status. Here, we conduct the first genome-wide association study (GWAS) on sociologically informed occupational status measures. We exploit our findings to advance sociological understanding and quantitative modeling of status attainment processes across the life course and their complex relationship with health.

The deeply engrained intergenerational transmission of SES and inequalities across generations^4,5^ has motivated social scientists to consider whether genetics plays a role in SES^6-8^ and, more recently, SES-related stratification, and non-genetic inheritance, which biases genetic effects on a phenotype.^9,10^ To date, the focus has been primarily on educational attainment^11,12^ and income,^13^ with less attention to the heritability of occupational status. However, family studies indicate moderate heritability of occupational status comparable to other SES measures in the range of 0.30 to 0.40.^14-18^ Molecular genetic research on SES proxies has focused on educational attainment^6,7,19-21^ and income,^22,23^ neglecting occupational status. SES measures have been recently highlighted as important in that they introduce gene-environment correlations which affect GWAS results^24^ and in influencing the patterns of genetic correlations of mental health traits.^25^ Hence, a more nuanced and complete understanding of SES that goes beyond educational attainment and income is needed. Whilst intertwined, these dimensions are clearly analytically and empirically distinct,^26^ and individuals may, for example, trade off income for other types of status, in particular occupations. Educational attainment may therefore not necessarily translate into economic success.

One GWAS on broadly skill-based occupational groups using the UK Biobank identified 30 genetic variants associated with 9 categories of the UK Standard Occupational Classification and a SNP-heritability of 0.085.^27^ However, occupation in the UK Biobank is richly measured using 353 categories and full information can be used to benefit from expert sociological theory and measurement of occupational stratification. In addition, purely skill-based measures have been criticized since they have inconsistent operationalization and are not comprehensively theorized, ignoring, for example, social prestige and other status factors.^1^

#### Box 1. Ethical Considerations of this Study

The study of genetics and its relationship with social status has a complex history, with some researchers in the past using biological factors to discriminate and reinforce inequalities. Early nineteenth-century work by Galton (1869)^28^ and other contemporaries linked biology to the study of intelligence, criminality, and status, leading to contentious debates on the validity and implications of their findings.^29-31^ Later studies, such as Sorokin’s (1927) “Social Mobility,”^32^ have been critiqued for assuming causality from correlations.^33^ Furthermore, works like “The Bell Curve” (1995) by Herrnstein and Murray revisited these earlier debates, suggesting a biological basis for societal stratification.

It is important to recognize that these past studies have contributed to an aversion and even fear of studying genetics in these areas of research.^29,34^ We recognize this apprehension and explicitly distance our research from studies that were overtly classist and/or racist and reinforced inequalities, confused structural inequality with biology or drew overly-simplistic policy implications. Our endeavor, rather, is rooted in the pursuit of a biosocial understanding of occupational stratification and the role that socio-economic status plays in genetic estimates, firmly guided by a well-established ethical and analytical framework.

In a 2023 consensus report from the Hastings Institute,^35^ a group of critical voices and researchers in the field of social and behavioral genomics emphasized the need for responsible conduct in studies examining the genetics of phenotypes deemed to be of “heightened concern,” particularly those with significant implications for social status. Although our analysis was conducted prior to the publication of this report, we note that our work already adhered to its study design guidelines.

Our investigation neither attempts to compare individuals across contentious social divisions in contemporary societies, such as race or ethnicity, nor does it involve comparing genetic ancestral groups that could be conflated with racial or ethnic categories. Our GWAS focuses solely on a population of British-European ancestry. We provide comprehensive justification for the definition and measurement of our key phenotypes and conducted our analysis using an adequately powered sample. Our results were replicated both out of sample and using within-family estimates, and we transparently discuss observed reductions in effect sizes. Although extending our study to other ancestral groups is beyond the scope of our current work, we encourage further research in this area.

We have included a frequently asked questions (FAQ) section in the Supplementary Material, which offers a clear and accessible explanation of what our findings do and, importantly – do not - find. Alongside the accurate scientific interpretation of our research, we advocate for open discussion on the importance of research into the genomics of socioeconomic status and the role that SES plays in patterning other genetic outcomes that engages scholars from a wide array of intellectual backgrounds and viewpoint diversity.

Sociologists consider occupation as the primary social and economic role held by most adults outside their immediate family or household, often even as “(…) the single most important dimension in social interaction” (p. 203).^36^ It is a long-term stable indicator of an individual’s social position in society alongside income, consumption, division of labor, and social reproduction as a whole.^1^ Adequately measuring occupational status is complex, with generations of sociologists dedicated to mapping this complex qualitative trait on a continuous scale.^37^ The three conceptual approaches to measuring occupational status either consider socioeconomic differences between occupations, inter-occupational social interaction, or ascribed prestige of different jobs.^38^

In our analyses, we focus on three different measures of occupational status, championed by different theoretical traditions in sociology. First, the International Socioeconomic Index (*ISEI),*^36^ is a status measure constructed from scaling weights that maximize the (indirect) influence of education on income through occupation. Second, the Standard International Occupational Prestige Scale (*SIOPS)*,^39^ is a prestige-based measure based on public opinion surveys where a representative population is tasked with ranking occupations by their relative social standing. The third Cambridge Social Interaction and Stratification Scale (*CAMSIS*),^1^ measures the distance between occupations based on the frequency of social interactions between them (operationalized as husband-and-wife combinations). This measure is based on the notion that differential association is a function of social stratification, with partners and friends more likely to be selected from within the same group. Although these measures are championed by different theoretical traditions in sociology, empirically they have substantial but not perfect correlations^40^ eluding to an underlying latent factor of occupational status.

The current study investigates molecular genetic associations with *ISEI, SIOPS,* and *CAMSIS*. Analyses were conducted on 273,157 (130,952 males and 142,205 females) individuals in the UK Biobank,^41^ identifying 106 genetic variants, and replicated in the UK’s National Child Development Study (NCDS; N= 4,899; 2,525 females and 2,374 males). Genomic structural equation modelling^42^ suggests a general genetic factor across all SES measures of occupational status, income, and education.

The integration of molecular genetics into this core topic of social science research promises a richer understanding of the role of the biological and the social as well as the improvement of quantitative modeling and understanding of social processes of attainment status transmission. We utilize our GWAS discovery results for various sociogenomic investigations in this regard. Whilst our understanding of the biological basis from GWAS findings for complex behavioral traits is still in its infancy,^19,43^ we investigated how social and psychological mechanisms play a role in the genetics of occupational status, including childhood career aspirations, noncognitive,^44^ and cognitive traits.^27^ We then examined to what extent polygenic scores (PGS) for occupational status predict the phenotype within and between families, their genetic penetrance of careers across the life course and the role common genetic variants play as a confounder of the intergenerational transmission of occupational status. Additional analyses explore the complex relationship between occupational status and health outcomes and how parental occupational status confounds the genetic prediction of general health. Our findings are conclusive that ignoring genetic data in parent-offspring SES transmission and quantitative stratification research in general leads to biased results in non-experimental studies, whilst the interplay between genes and the environment remains complex.

## Results

### Heritability, discovery, and genetic relationship amongst SES measures

The main analyses was conducted on individuals from the UK Biobank on the three phenotypic measures of occupational status: CAMSIS (N=273,157), SIOPS (N=271,769), and ISEI (N= 271,769; see Methods). Linkage Disequilibrium Score Regression (LDSC) based SNP-heritability (ℎ^2^_SNP_)^45^ was significantly different from zero for all occupational measures, whilst ∼50% larger for CAMSIS (ℎ^2^_SNP_= 0.145, SE = 0.0066) compared to SIOPS (ℎ^2^_SNP_= 0.105, SE = 0.0052) and ISEI (ℎ^2^_SNP_ = 0.109, SE = 0.0056, see Figure 1). This is within the range of ℎ^2^_SNP_ for other status measures estimated in the UK Biobank (see Methods) such as education (ℎ^2^_SNP_ = 0.153, SE = 0.0056) and income (ℎ^2^_SNP_ = 0.092, SE = 0.0041) and for CAMSIS nearly twice as high as for previously reported occupational measures.^27^

**Figure 1.**
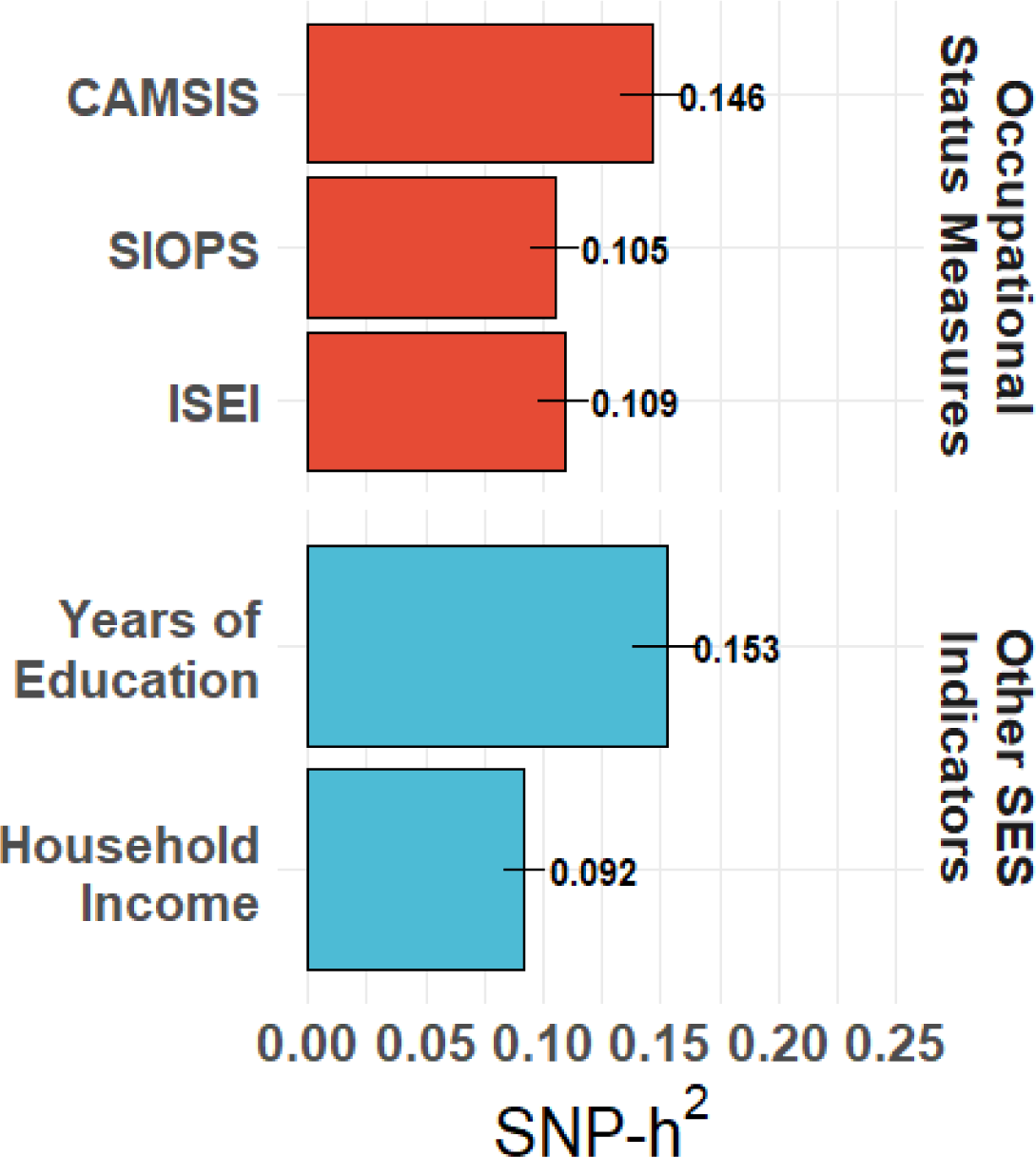
LD score-based SNP-heritability of occupational status measures CAMSIS (N = 273,157), SIOPS (N = 271,769), and ISEI (N = 271,769) on top compared to income (N = 353,673) and education (N = 404,420). Bars denote 95% confidence intervals.

The GWASs identified 106 genetic variants for CAMSIS including 56 also found for ISEI and 51 for SIOPS based on an *R*^2^-threshold of 0.1 and a window-size of 1000kb (see Figure 2 for the Manhattan plot), one of which (only significant for CAMSIS) was found on the X-chromosome. We identified 11,206 SNPs in LD with our autosomal lead SNPs (see Methods) and conducted an exhaustive phenome-wide association study (PheWAS) using the GWAS catalogue and the IEU Open GWAS Project database. While we observe a substantial overlap with other socioeconomic status-related traits, 8 of our variants (rs12137794, rs17498867, rs10172968, rs7670291, rs26955, rs2279686, rs72744938, rs62058104) have not yet been linked to any status-related trait. For three variants (rs7670291, rs26955, rs72744938) not even suggestive associations (p < 5×10^-6) with status traits are discernible. For two of these, we find strong links to platelet count. A full list of all implicated phenotypes is provided in Supplementary Table B7. The only non-autosomal hit (rs146852038) has previously been linked to the age of first sexual intercourse and educational attainment.

**Figure 2.**
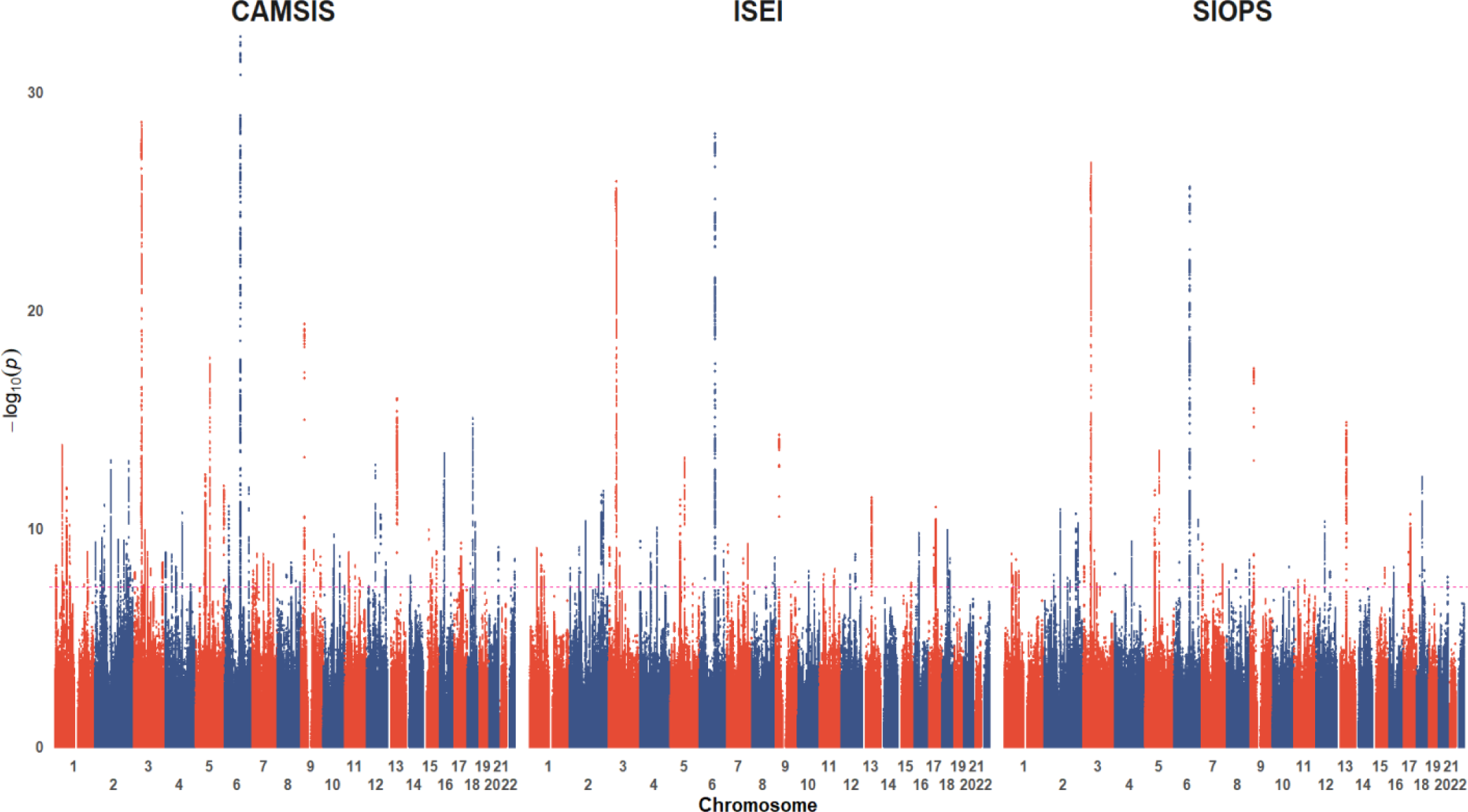
Manhattan Plot with autosomal SNP position on the X-Axis and the logarithm of the p-value on the Y-Axis of the GWASs for Occupational Status Measures, from left: CAMSIS (N = 273,157), ISEI (N = 271,769) and SIOPS (N = 271,769).

We then replicated these hits using the National Child Development Study (NCDS), an ongoing study of a British birth cohort born in 1958 (see Methods). This dataset was chosen since it is a similar UK cohort, given that previous research has demonstrated variation by country and birth cohort, particularly for complex behavioral phenotypes.^46^ Despite the notable disparity in sample size, with 4,899 individuals in the NCDS compared to ∼273,157 in our discovery sample, our results surpassed the expected sign concordance and achieve a higher than anticipated number of significant hits at *p* = 0.05 (see SI 6.4). This replication result underscores the robustness of our findings, even when subjected to the constraints of a smaller sample.

To investigate the functional implications of the genetic variants associated with occupational status, we performed gene-based and gene-set analyses using MAGMA (see Methods).^47^ We observe that genes implicated by our SNPs are expressed in the brain, including the pituitary gland. No other tissue showed significant enrichment for gene expression.

We also jointly analyzed the highly correlated occupational status measures together with income and education to increase statistical power using multi-trait analysis of GWAS^48^ (MTAG, see Methods) resulting in 731, 646, and 653 variants passing the significance threshold for CAMSIS, ISEI, and SIOPS respectively.

Genetic correlations (Figure 3, lower left triangle) between the three measures were close to 1, and thus stronger than the phenotypic correlations (upper right triangle), ranging between 0.80-0.90. The genetic correlations with educational attainment and household income were almost twice as high (0.81-0.97) as their phenotypic counterparts (0.32-0.44). Considering these high genetic correlations, it is unsurprising that we found strong evidence for a common genetic factor of occupational status using Genomic Structural Equation Modelling (GSEM)^42^ with high loadings for all three measures (standardized path coefficients of 0.99, 0.99, and 0.99, for CAMSIS, ISEI, and SIOPS, respectively; see Supplementary Information Section 9). We furthermore provide evidence for a common genetic factor of SES including income and education (see Supplementary Figure 6).

**Figure 3.**
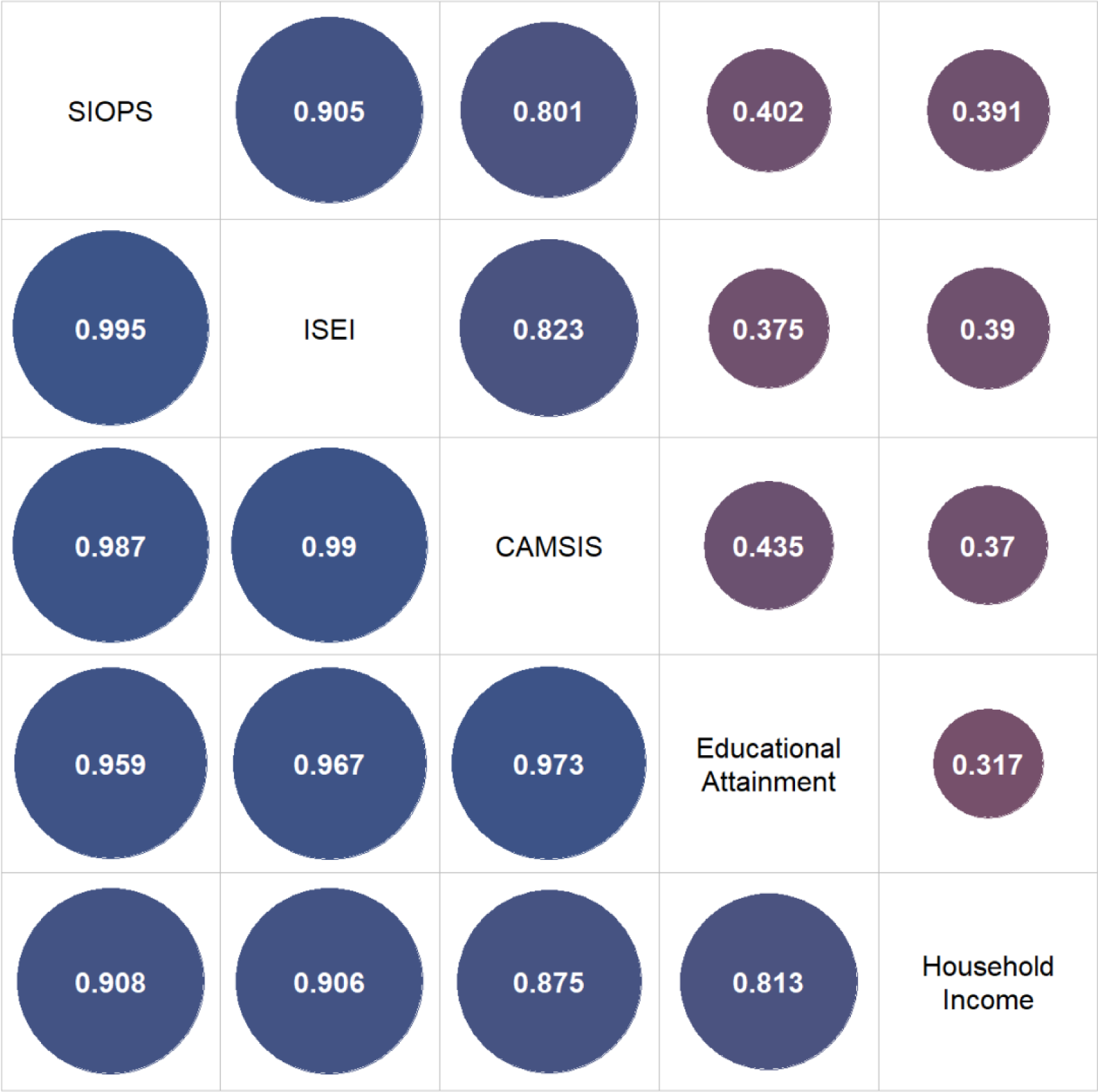
Phenotypical (upper right triangle) and genetic correlations (lower left triangle) of occupational status measures and other SES indicators based on LD score regression. N for phenotypical correlations = 246,492.

### Polygenic prediction

We assessed the out-of-sample predictive performance of the PGSs using two data sources. The first sample comprised a subset of siblings from the UK Biobank, for which we conducted an additional GWAS while excluding these individuals from the discovery analysis. The second sample consisted of the aforementioned NCDS.

MTAG-based out-of-sample predictions were nearly identical in both UK data sets, with an incremental *R*^2^of 0.077 (SE=0.00306) in NCDS across all observations (0.075, SE=0.00287 in the UK Biobank) for CAMSIS, 0.054 (SE 0.00306; 0.054, SE 0.00250) for ISEI and 0.055 (SE=0.00255; 0.053, SE = 0.00248) for SIOPS (Figure 4).

**Figure 4.**
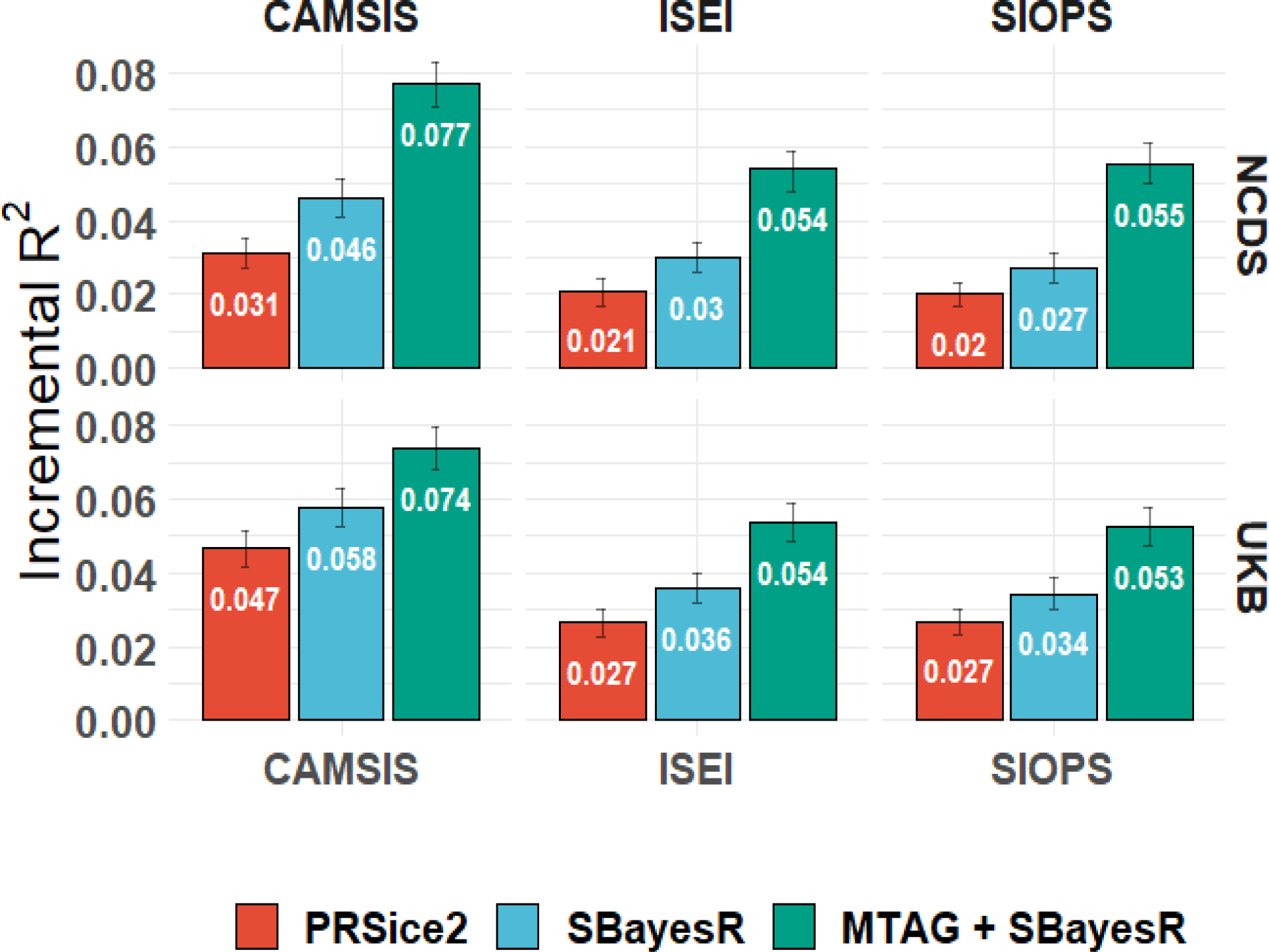
Out of sample polygenic prediction performance within UK Biobank and NCDS. Incremental *R*^2^ compared to a baseline model consisting of 10 principal components, sex, and age. Bars denote 95% confidence intervals. N = 24,579 for CAMSIS and 24,472 for ISEI and SIOPS in the UK Biobank, for NCDS average performance over different ages (N = 5,389; 5,312; 5,211; 4,902; 4,263 for CAMSIS at age 33, 42, 46, 50 and 55, N = 5,449; 5,293; 5,197; 4,892; 4,252 for ISEI/SIOPS)

The longitudinal data in the NCDS reveal changes in the PGS effects across the life course or career trajectories, respectively. First, we were able to examine PGS prediction of occupational status across the life course, at ages 33, 42, 46, 50, and 55, noticing slight heterogeneities, with the highest predictive accuracy achieved at the beginning and end of the career with more than 9% and the lowest of 3% measured at age 42 (see SI 11).

We then ran growth curve models to estimate the slope of PGS effects of occupation and its quadratic term. Trajectories vary significantly by the polygenic signal for all three measures: The initial genetic stratification effects are compressed mid-career and re-establish towards the end of the career (see Figure 6). At the lower end of the PGS distribution, status at the beginning of the career (age 33) is comparatively low but increases steeply for roughly a decade before after either stabilizing or even decreasing again. At the upper end of the distribution, however, a much higher starting prestige is followed by accumulation or compounding status gains, which show no sign of stopping up to age 55. Further analyses indicated that for CAMSIS, these findings were robust to the inclusion of controls for parental SES in the model, while for SIOPS neither parental SES nor the PGS was significant when jointly modeled (See Supplementary Information 14).

**Figure 5.**
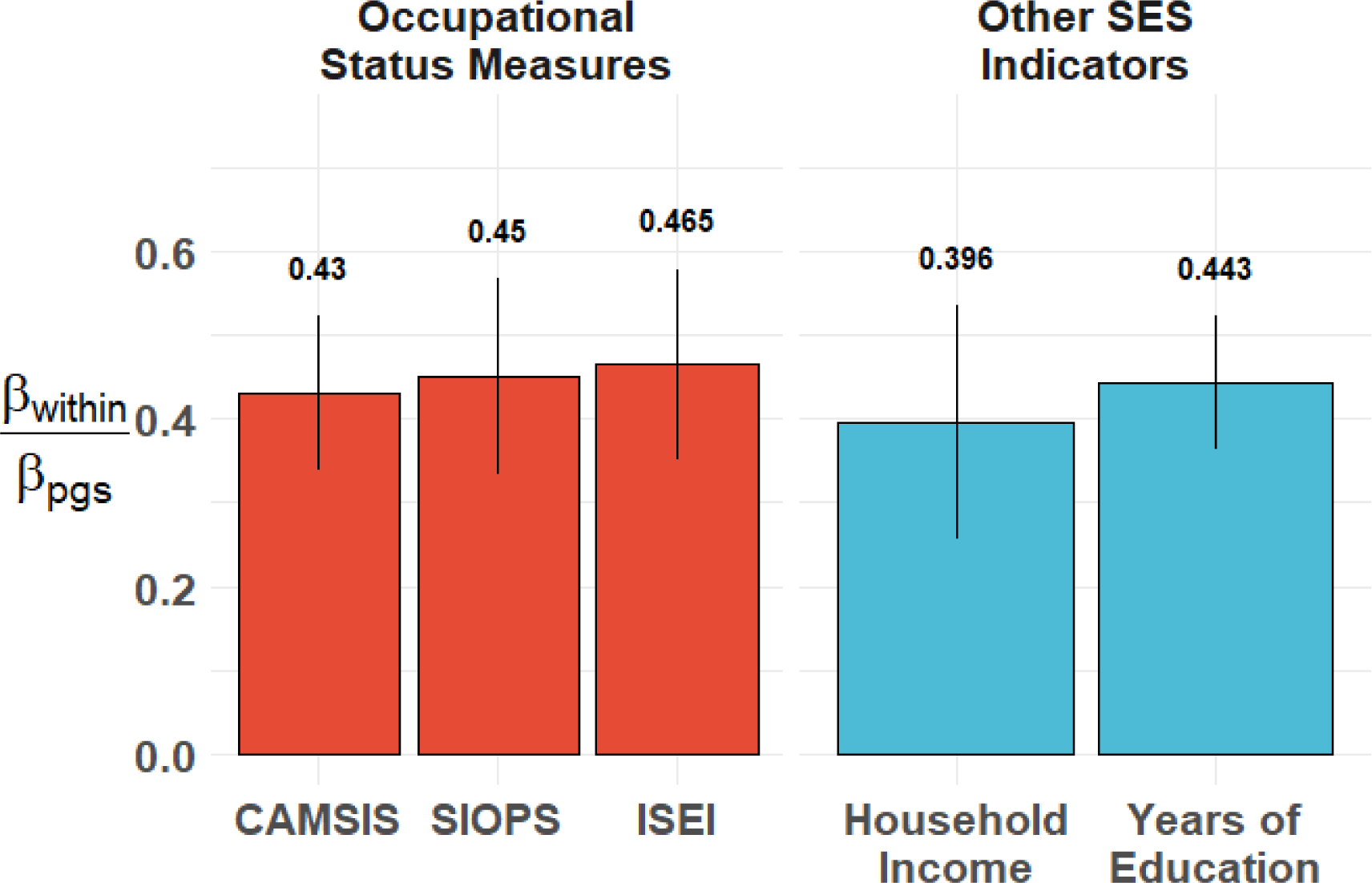
The ratio of standardized beta coefficients for effect of the respective PGS on the phenotype based on between and within family models, for CAMSIS, SIOPS, ISEI, household income, and years of education among siblings in UK Biobank. N = 24,579 for CAMSIS, 24,472 for ISEI and SIOPS, 36,265 for education, and 31,851 for income. Bars denote bootstrapped 95% confidence intervals.

**Figure 6.**
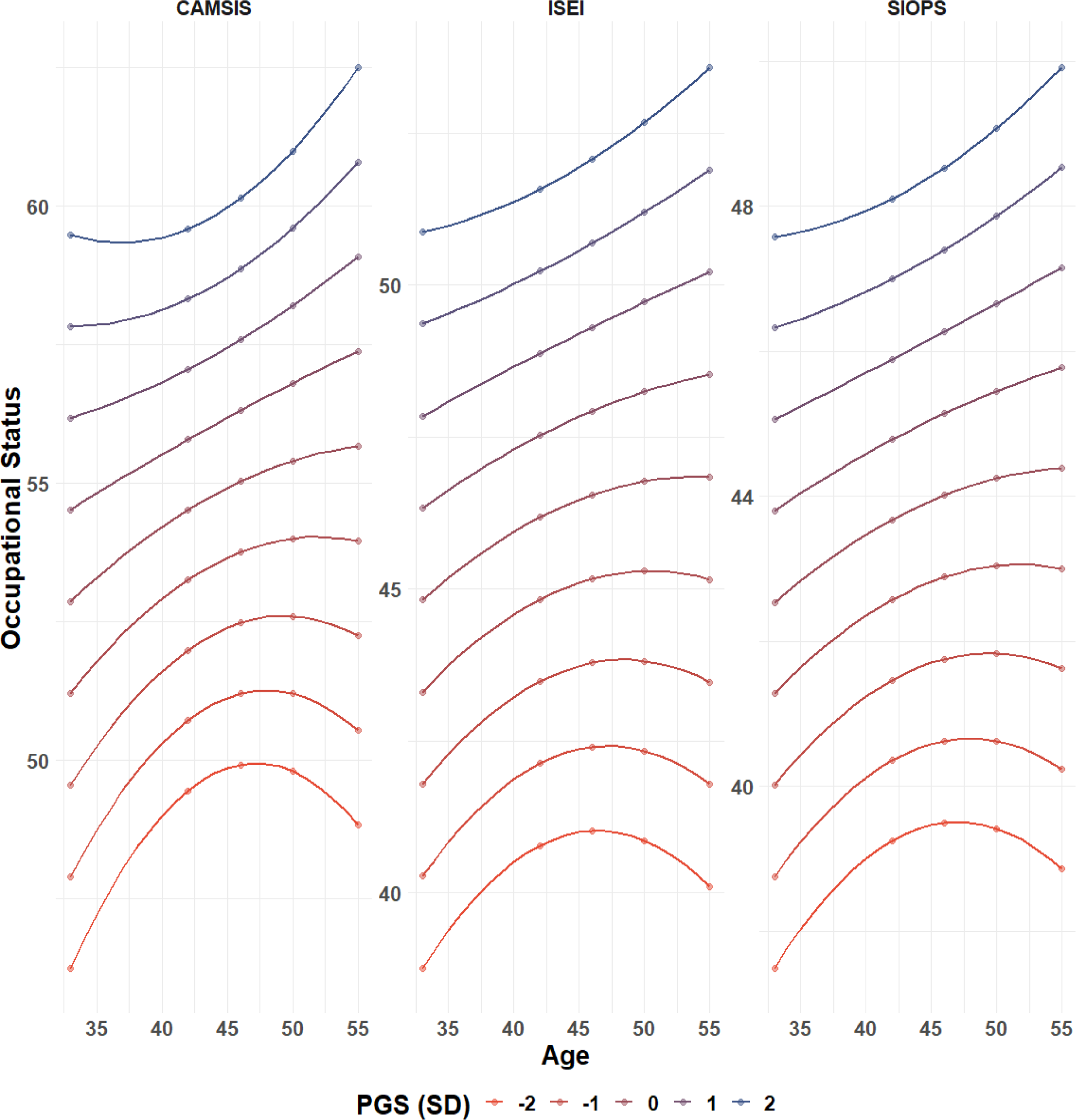
Growth curve models with varying slopes depicting the predicted values for the three occupation measures across the life course between ages 33 and 55 over the whole PGS distribution (-2 SD to +2 SD). Dots mark observed years. N = 2,883 (CAMSIS); N = 2,894 (SIOPS/ISEI)

### Disentangling direct, indirect, and demographic effects

GWAS population estimates include a combination of direct effects (inherited genetic variation), and indirect effects or gene-environment correlations and can be further influenced by assortative mating. We conducted multiple analyses to better understand the relative importance of these dimensions in relation to our estimates (see Supplementary Information 8 & 10).^49,50^

First, we investigated the predictive performance of our scores between more than 29,500 siblings in the UK Biobank, a common design to identify direct genetic effects. Notably, traits related to socioeconomic status or other non-clinical outcomes tend to exhibit considerable within-family effect reductions^38,41^, potentially affecting their practical utility^51^. Our analysis supports these previous studies showing a reduction in effects for occupational status measures of more than 50% in total, with results for other SES measures (education and income) in a similar range (see Figure 5 for the ratio of population and within-family models and Methods).

This discrepancy between the unrelated population and within-family estimate can be attributed to indirect family effects or assortative mating. Indirect effects include the (heritable) social transmission of economic resources, and cultural and social capital, as well as social-psychological factors such as parental expectations, which represent passive gene-environment correlation. To quantify the role of indirect effects, we follow two research designs. First, we adjust the best-performing PGS prediction in the NCDS for parental SES (measured as parental occupational status at age 11). Second, we conduct an adoption prediction study. In an adoption design, children are raised by non-biological parents, thereby providing a unique opportunity to examine the influence of genetic factors while minimizing the effects of passive gene-environment correlation. We re-ran our GWAS for occupational status, while excluding the set of 3,414 respondents of European ancestry in the UK Biobank that stated that they were adopted and for which occupational information was available. Results from both designs are remarkably similar with the parental SES control design reducing the incremental PGS *R*^2^to 0.044, 0.031, 0.028 for CAMSIS, SIOPS, and ISEI, with an effect attrition of 22%, 22%, and 22%, respectively, and the adoptee prediction resulting in an *R*^2^ of 0.043, 0.031, 0.027 for CAMSIS, ISEI, SIOPS respectively which reduces to an effect reduction of 23%, 22%, and 27%.

The observed remaining discrepancy between the population estimate controlling for indirect effects and within family estimates could be attributed to attrition in the within family design due to assortative mating, which attenuates the within-family effect. Recent findings by economic historians have demonstrated strong partner matching on occupational status within the UK dating back to at least the 1750s.^52^ By employing a method first proposed by Lee et. al.,^6^ we demonstrate that, even in the absence of indirect effects, within-family effects are plausibly anticipated to be attenuated by 22-27% (SI 10), which closes the observed gap between both estimates. Under the assumption of additive effect reduction due to assortative mating and indirect effects, all three methods consistently estimate the proportion of direct population effects to be within the range of 73-78%. This convergence of findings underscores the importance of accounting for biases related to partner matching when examining the role of genetic factors in occupational status. It furthermore motivates the inclusion of parental SES for robustness in the application of PGS analyses downstream the population GWAS.

### Social mechanisms linking genetics and occupational status

A pertinent question to consider is which traits serve as mediators between an individual’s genome and occupational status. Evidence from twin studies indicates that both cognitive and noncognitive traits play a mediating role in the relationship between genetic and social outcomes.^53^

Building on previous behavioral phenotype GWASs and the literature,^54^ we identified five causally upstream traits that are potential mediators of the general genetic factor of SES: cognitive performance,^6^ ADHD as a proxy for behavioral disinhibition,^55^ openness to experience,^56^ risk tolerance,^57^ and neuroticism.^58^ In a multivariate genetic regression model (see Supplementary Information 9.3), overall, we can explain 70% of the genetic association with occupational status, but cannot, however, disentangle the unique contributions of each factor to this explanation.

In the NCDS data, we, therefore, tested the mediating effects for the occupational status PGS of the phenotypic measures of cognitive ability, externalizing behavior, internalizing behavior, scholastic motivation, and occupational aspiration at age 11 (see Methods and SI 13). Depending on the career stage of the respondents indicated by NCDS waves, these variables explained 54-74% of the link between our PGSs and occupational status. As expected, cognitive ability was the main mediator, explaining 31-50% of the association depending on respondents’ age. Scholastic motivation explained between 7-12%, occupational aspiration 6-11%, and other non-cognitive traits only up to 5%. Effect reductions are proportional when adjusting for parental SES in order to control for passive gene-environment correlation or indirect effects respectively (Supplementary Information Section 13).

### Intergenerational transmission

Given that parental status is strongly associated with their offspring’s status, the study of intergenerational status transmission has a long tradition, often focusing on educational attainment.^59,60^ In the NCDS data, the phenotypic correlation between paternal occupation at age 11 and offspring occupational status at various ages for all three measures was substantial (∼.30). Including a PGS to control for genetic inheritance and identify social effects reduced the intergenerational correlation of occupational status by 11%. However, this is likely an underestimation given the power limitations of GWAS in capturing full SNP-heritability.

Rescaling the results to estimated SNP-heritability^61^, ∼38% of the intergenerational correlation is due to common genetic inheritance and ∼62% is due to social inheritance and possibly the effects of rare genetic variants^62^ not captured by SNP-heritability estimates (see also Figure 7 and Supplementary Information 12 for estimates by age).

**Figure 7.**
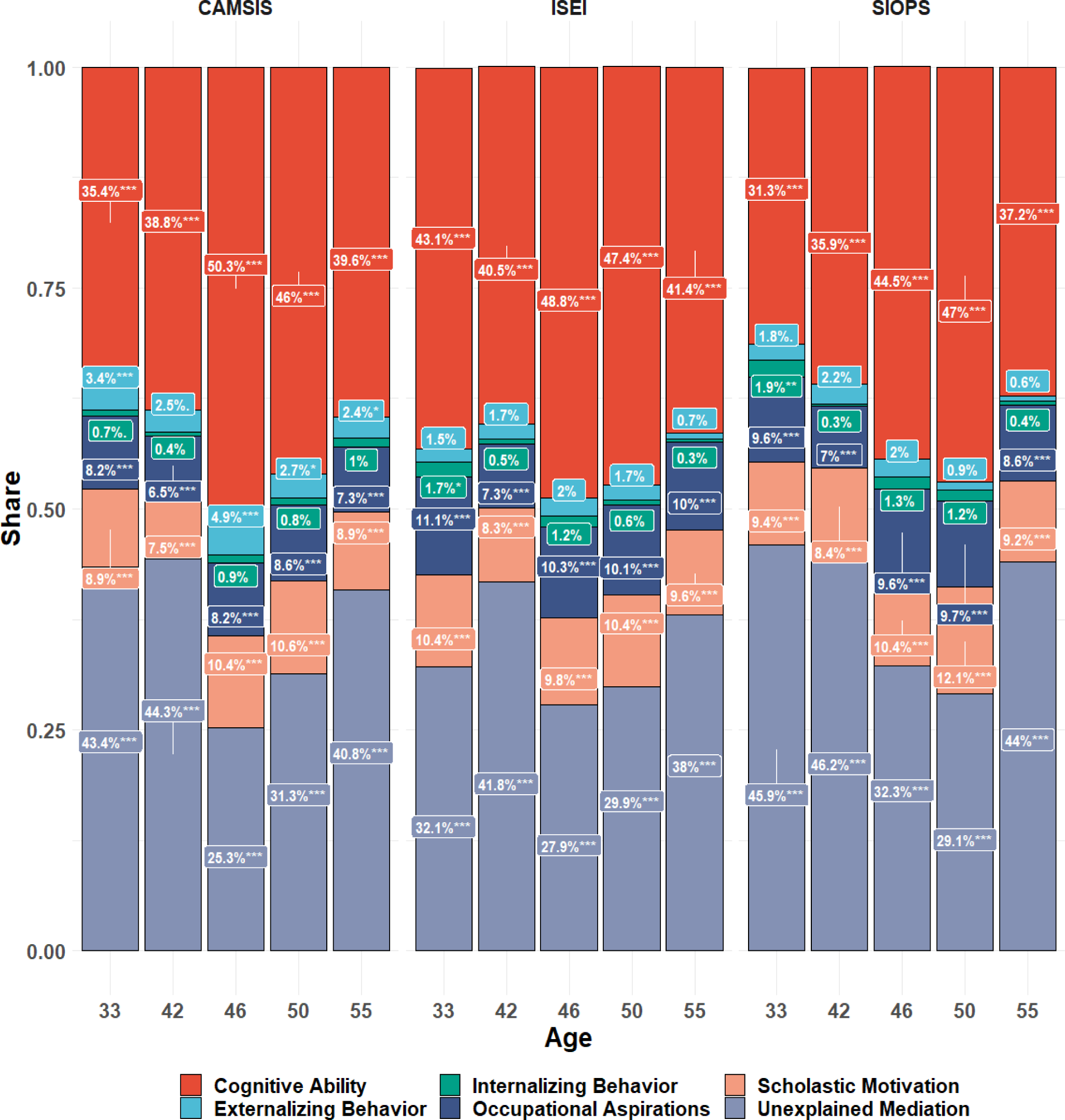
Depicting the percentage of polygenic score effects on occupational status mediated by cognitive and non-cognitive traits in NCDS through the life course. N = 3,169; 3,111; 3,075; 2,881; 2,499 for CAMSIS at age 33, 42, 46, 50, 55 and 3,196; 3,100; 3,068; 2,878; 2,494 for SIOPS/ISEI. Significance levels based on p-values are represented as *** for p < 0.001, ** for 0.001 ≤ p < 0.01, and * for 0.01 ≤ p < 0.05.

### Genetic confounding in the relationship between occupational status and health

Occupational status is correlated with various health outcomes and higher SES individuals typically live longer and in better health.^2^ It is essential to understand to what extent this association between occupational status and health is a causal one in order to, for example, design effective intervention strategies. The observed association could partly be driven by endogeneity since individuals in better health also potentially secure better jobs or perform better at work. Controlling for genetic associations reduces biases arising from genetic endogeneity also in regards to potential direct pleiotropic effects.^63^ We, therefore, investigate to what extent the occupational status PGS confounds the observed relationship between occupational status and general health as well as mental health in the NCDS data (see Supplementary Tables 7 & 9 for regression estimates). Similar to the intergenerational transmission of status, we find significant genetic confounding in the observed relationship. The association reduces on average across ages and outcomes, with 29% predicting general health and 13% predicting mental health.

To better understand the degree to which the genotypic effect of occupational status on general and mental health might incorporate indirect effects, we analyzed the health of the respondents based on their occupational status PGSs with and without parental occupational status at age 11 as a control variable. In accordance with previous results, we found that taking parental occupational status into account reduced the PGS prediction of general health on average across ages and outcomes by 19.5% and mental health by 23.7%, demonstrating the importance of considering parental SES indicators for the genetic study of offspring’s health outcomes (see Supplementary Tables 8 & 10).

## Discussion

Analyzing data from 273,157 individuals from the UK Biobank, we identified 106 genetic variants associated with occupational status measures, 8 of which have not been previously reported in related SES GWAS. Our study provides PGSs that measure the genetic underpinnings of occupational status, with an out-of-sample prediction of 5-8% depending on the status measure and up to 9% depending on the career stage. Genetic effects derived from the CAMSIS were around 50% larger than for SIOPS and ISEI and twice as high as naïve measures applied in the life sciences.^27^ We, thus, provide the R-package ‘*ukbjobs*’ to equip all researchers working with the UK Biobank to use these well-defined sociologically informed measures.^64^ CAMSIS conceptually focuses on social interactions – in contrast to, for example, purely skill-based measures. A potential reason for this observation may be genetic selection into interaction networks^65,66^. However, high genetic correlations between CAMSIS, SIOPS, and ISEI might also point to the benefits of a more granular and exact measure of the same latent phenotype in CAMSIS.

Our study not only demonstrates the genetic interdependence of occupational status measures, but also unveils a very strong genetic correlation between educational attainment, income, and occupational status, identifying a common genetic factor of SES. Notably, genetic correlations amongst SES indicators surpass phenotypic correlations by a factor of two to three. This outcome represents an outlier from Cheverud’s (1988) conjecture,^67^ which states that phenotypic correlations can serve as proxies for genetic correlations—a notion that finds empirical support in both animals and humans.^68,69^

The deviation might have several reasons including trade-offs between investments into different dimensions of SES. Higher education does not always guarantee high income or occupational status, since labor market conditions, personal networks, and ethnicity and gender can influence career trajectories.^5^ Likewise, higher occupational status does not always bring a high income or require advanced education, and the importance of occupational status may vary across cultures and social contexts.^70^ Certain genetic traits may predispose individuals to achieve higher levels in these areas through biological pleiotropy.^23^ For instance, genetic factors influence cognitive abilities, personality traits, and mental health, which may, in turn, impact educational attainment, income potential, and occupational status. However, environmental factors such as family background, social norms, cultural values, societal expectations and chance are also highly influential in shaping education, income, and occupational status. Environmental differences in individual cases can lead to more heterogeneity and thus weaker phenotypic correlations and subsequently have a completely different causal pathway in influencing health and behavioral outcomes.^26^

In a general framework of social mobility, economists proposed that historically variable measures of socioeconomic status are transmitted via a potentially genetically influenced latent factor.^4,71^ Occupational status, income, and education might currently operate within the British population as heterogeneous phenotypic manifestations of such a latent general genetic factor of socioeconomic status.

We have extensively shown that the prediction attrition within families is in part due to indirect genetic effects or genetic nurture respectively, which also consistently contribute to the latent factor for constructed SES measures. Moreover, a mounting body of evidence suggests that strong assortative mating on this latent factor that has been present for multiple generations.^52,72^ Notably, a higher spousal correlation has been observed for the genetic predictor of educational attainment than for the actual phenotype.^73,74^ This phenomenon may partially account for why causal genetic variants display a stronger predictive power for occupational status between families, as opposed to within families where the variation in these variants is more limited.

We integrated the polygenic signal for occupational status into occupational mobility and social stratification research and vice versa as it has crucial implications on both sides. First, intergenerational mobility in social status is of great interest, not only for social scientists such as sociologists and economists, but also policymakers, public health and epidemiology and is related to questions of equality of opportunity, and commonly used to indicate societies’ degree of openness.^5,75^ The role of genetics has been interpreted as a cause of intergenerational status resemblance independent of social inequalities, indicating merit. We find evidence that next to cognitive skills, also scholastic motivation, occupational aspiration, personality traits, and behavioral disinhibition (proxied by ADHD), drive the association between genetics and occupational status. Importantly, the mediating factors are highly polygenic and contingent on the population in which they emanate. It is also vital to note that around one-third of the polygenic signal remains unexplained in each of our approaches. This urges the need for further investigations to better understand the role of genetics in status inheritance and to comprehensively evaluate the interpretation of heritability as a pure merit measure in the context of questions addressing equality of opportunity.

Second, when examining the genetics related to the intergenerational transmission of SES, mostly heritability studies on educational attainment assumed that genetic influences are stable in absolute terms and environmentally driven inequality reduces with lower intergenerational correlations.^8^ Extrapolating results from PGSs, we show that the intergenerational correlation for occupational status is up to 38% due to genetic inheritance – this is even stronger than for educational attainment ^59,60^. This suggests that social stratification researchers need to adjust their sole focus on intergenerational correlations to also explicitly consider gene-environment correlation in their statistical modeling. We note that the applied extrapolation assumes SNP-heritability levels but could still represent an underestimation since PGSs have a lower prediction compared to SNP-heritability, but the latter is still smaller than the heritability estimated from twin models. The discrepancy between SNP- and twin-heritability might be due to rare genetic variants, higher environmental homogeneity within families, and non-linear genetic effects.^46,76^ As often noted, however, twin studies likely also overestimate heritability due to the violation of the assumption that identical and fraternal twins share environmental influences to the same extent. SNP-heritability as measured here remains a conservative approach compared to previous studies.^59^

Third, we highlight questions about the causality of the relationship between health and occupational status and SES in general.^2^ It is plausible to assume that higher status causally leads to better health, for example, due to a higher living standard, nutrition, and better knowledge about and access to health care systems, amongst others.^2^ At the same time (heritable) poor health might also select an individual into lower-status occupations, or genetics might have direct pleiotropic effects on education and health or related factors, leading to an overestimation of a direct, phenotypic causal effect. The question of causality, however, is paramount for designing targeted policy interventions and genetic confounding needs to be considered. It is also relevant to quantify their potential impact and clarify claims in social mobility research regarding genetically driven, health-related confounders. We show that the association between occupational status and health is up to 45% confounded by genetic effects and therefore substantially upward-biased, when genetic factors are not considered.

Fourth, combining theoretical, measurement and modelling perspectives of the social sciences and genetics is not only important for the interpretation of status in social science theory and modeling, but also for genetic research.^76^ First, the discovery of genetic nurturing effects unraveled the importance of social influences correlated with genotypes in the discovery of genetic effects on education.^9^ We show that controlling for parental occupational status strongly reduces genetic prediction of the occupational status PGS with general and mental health. While genetic prediction based on our PGSs on health is comparably small (1%) and confounding effects may not entirely generalize to other regions of the genome important for health outcomes, further investigation is required to understand whether and how parental SES measures should be integrated in population GWAS studies where available. Second, the continued use of expertly developed measures that have a strong theoretical, conceptual, and measurement basis such as occupational prestige in social stratification research, underlines the importance of precision phenotypes. Contrary to a prior GWAS that relied solely on a skill-based minimal occupational classification,^27^ our occupational prestige phenotypes, which have been meticulously developed and refined by sociologists over decades, doubled the heritability using CAMSIS, increased SNP discovery by more than threefold and also provides a consistently meaningful interpretation of the outcomes variable. This also emphasizes the genetic relevance of socially theorized measures and of social factors included in them, such as potential interaction or social prestige.

Finally, our findings embrace a comprehensive and interdisciplinary perspective when studying social stratification, mobility, and status transmission. By further studying the underlying latent factor of individual socioeconomic status indicators, we can foster a more profound understanding of the genetic basis of socioeconomic status and its broader implications for society. It is imperative to comprehend the role of indirect effects and passive gene-environment correlations in this puzzle as well as the causes and consequences of assortative mating on these relationships. The dynamic nature of the intergenerational transmission of socioeconomic status is most prominently synthesized across social, historical, and genetic theories in the future applied to data into a comprehensive and rigorous model.

Our study is also not without limitations. The UK Biobank data represent only 5.5% of the approached target population and is highly selective in several ways, overrepresenting individuals with lower genetic risk of mental health problems, BMI, nonsmoking, higher education, and from less economically deprived areas^49,77-80^ and alternative data sources would be useful for discovery and prediction. In this context, we can expect environmental heterogeneity across different populations to challenge our findings. While PGS predictions are nearly identical in our two UK populations, previous research has demonstrated that for educational attainment, only 50% of genetic effects are universal across seven Western populations.^46^ Population genetic heterogeneity also limits the scope of this study beyond UK residents, and as the majority of GWAS to date, we only focus on European-ancestry individuals in a Western country.^81^ The integration of other ancestries, geographical and more diverse socioeconomic contexts is the future. The reduction of PGS prediction within families, impacting predictability and potentially other analyses, also emphasizes the relevance of recent initiatives for discovery designs using family data and to further study the role of assortative mating for within-family effect reduction.^49^ Despite these limitations, the current study offers many new insights into the genomics of occupational prestige and socioeconomic status.

## Methods

This article has Supplementary Information (SI) with details about data and methods and additional detailed analyses. We have also built an R-package ‘*ukbjobs*’ that allows researchers to construct CAMSIS, ISEI, and SIOPS occupational scores directly from the UK Biobank data (https://github.com/tobiaswolfram/ukbjobs).

### Samples

### UK Biobank

For both, the discovery and prediction of occupational status measures, education, and income, we used data from the UK Biobank. The UK Biobank is a large-scale biomedical database and research resource, containing in-depth genetic and health information from 502,655 individuals recruited between 2006 and 2010. The database is globally accessible to approved researchers undertaking vital research. Ethical approval for UK Biobank was received from the Research Ethics Committee of the Department of Sociology (DREC) at the University of Oxford (SOC_R2_001_C1A_21_60). This work was conducted under UK Biobank application 32696. Details of the UK Biobank genotyping procedure can be found elsewhere.^82^ After phenotype selection and genetic Quality Control, we conducted our analyses on 273,157 individuals (130,952 males, 142,205 females)

### NCDS

As a second, longitudinal UK prediction sample from the UK, we used The National Child Development Study (NCDS) following 17,000 children born in Great Britain in one week in 1958 (application number GDAC_2021_16_TROPF). It has been designed to examine the social and obstetric factors associated with stillbirth and death in early infancy. Overall, there are ten waves available (birth – 1958, age 7 – 1965, age 11 – 1969, age 16 – 1974, age 23 – 1981, age 33 – 1991, age 42 – 2000, age 46 – 2004, age 50 – 2008 and age 55 – 2013).

### Phenotyping

S*ocioeconomic differences*-based indices measure the “attributes of occupations that convert a person’s main resource (education) into a person’s main reward (income)”.^36^ The most commonly used measure is occupational prestige, termed the *International Socioeconomic Index* (ISEI),^36^ which is constructed from scaling weights that maximize the (indirect) influence of education on income through occupation.

Other prestige-based measures are the result of public opinion surveys in which representative samples of the population are tasked with ranking occupations by their relative social standing. Emerging at a similar time as socioeconomic indices,^83^ Treiman (2013)^84^ demonstrated that prestige-based measures were surprisingly constant over time and cultures, consolidating their use in social scientific research. The *Standard International Occupational Prestige Scale* (SIOPS or Treiman-prestige)^39^ remains one of the most commonly used metric in this tradition.

Lastly, occupational status indicators derived from *social interaction* focus on the heterogeneity of associations between occupants of different jobs, following the tradition of Warner, Meeker, and Eells (1949)^85^ and Laumann and Guttman (1966).^86^ They are based on the idea that differential association is a function of social stratification since members of a group are more likely to interact within that group than with out-group members. Thus, acquaintances, friends, and spouses are much more likely to be selected from within the same group than from outside. A group of Cambridge sociologists reversed this approach to measure social structure based on interactions. The *Cambridge Social Interaction and Stratification Scale* (CAMSIS) measures the distance between occupations based on the frequency of social interactions (operationalized as husband-and-wife combinations) between them.^1^

Information on occupational status scales was merged to the occupational classification scheme utilized in UKB (SOC2000).^87^ *CAMSIS*-based status could be directly merged using the data available at Lambert and Prandy (2018).^88^ *ISEI* and *SIOPS* (as provided by the R-package “strat”),^89^ however, use the less granular ISCO-88 scale, so a mapping from ISCO to SOC was employed.^90^ If multiple job codes for a respondent were available, the most recent job was used.

*Income* was measured similarly as in Hill et al. (2019)^23^ using a coarse, 5-level ordinal household income variable. Educational attainment was defined as years of education and coded according to the scheme provided by Lee et al. (2018).^6^

The prestige of *current* or *most recent occupation* is treated as a continuous measure. In the initial discovery analysis using the UK Biobank, respondents were asked to provide job titles for the current or the most recent job held. The job information was coded using the four-digit UK Standard Occupational Code version 2000 (SOC2000). We built a procedure to link the UK SOC2000 to ISCO-88(COM), and then derive ISEI and SIOPS from ISCO-88(COM). All phenotypes are inverse-normal rank transformed before analysis. In the NCDS, the SOC2000 code of the respondent’s occupation (as well as their father’s when they were 11 years old), is available as well, so the same procedure was applied.

In the NCDS, we look at *health* measured at ages 23, 33, 42, 46, 50, and 55. Participants were asked to rate their general health on a scale from one (excellent) to four (poor) (age 23 and 33), one (excellent) to five (very poor) (age 42), one (excellent) to five (poor) (age 46, 50 and 55).

For each time point, the outcome is treated as metric and standardized to have a mean of zero and a standard deviation of 1. We then regress it on the CAMSIS, ISEI, and SIOPS PGSs, respectively, while controlling for sex and ten principal components to correct for population stratification.

### Discovery

An analysis plan was pre-registered and uploaded by Brazel, Ding, and Mills on February 2021 (https://osf.io/329pr/) and updated in February 2023 (https://osf.io/x6va5). All calculations are based on mixed model association tests as implemented in the program FastGWA,^91^ with association testing based on v3 imputed data. Following the pre-posted open science analysis plan in each regression, the following covariates are included: The first 10 genomic principal components, age at assessment and age^2, UK Biobank assessment center at recruitment, sex, genotyping array (BiLEVE or Axiom) on the sample of European ancestry. Chromosomes are analyzed separately. To speed up the calculation of summary statistics, a minimum MAF filter of 0.01 was imposed, leaving 10.2 million SNPs for the analysis. We supplemented our autosomal analyses with association analyses of SNPs on the X chromosome in a joint association analysis of both sexes.

### PheWAS

All 1000 Genome SNPs in linkage disequilibrium (*R*^2^> 0.6 for European ancestry) with the 106 were identified using FUMA.^92^ For these 11,206 SNPs, 1,005,470 phenotypical associations reaching at least suggestive significance (p <5×10^-6) in the GWAS catalogue and the IEU OpenGWAS project were collected.^93^ All variants with at least one genome-wide significant link to a trait associated with education, income, or any other socioeconomic outcome were removed, leaving 8 hits (rs12137794, rs17498867, rs10172968, rs7670291, rs26955, rs2279686, rs72744938, rs62058104) that have not yet been linked to any status-related trait. For three variants (rs7670291, rs26955, rs72744938) not even suggestive associations (p > 5×10^-6) are found.

### Univariate LDSC

Univariate LDSC regression was performed on the summary statistics from the GWAS in order to quantify the degree to which population stratification influenced the results and to estimate heritability. For this, GWA test statistics were regressed onto the LD score of each SNP. LD scores were used with European populations and weights were downloaded from https://utexas.app.box.com/s/vkd36n197m8klbaio3yzoxsee6sxo11v. SNPs were included if they had an MAF of >0.01 and an imputation quality score of > 0.9 and were available in the LD score file. Intercepts for all three occupational status measures were close to 1 (CAMSIS: 1.1193, SE = 0.013, SIOPS: 1.0845, SE = 0.011, ISEI: 1.0993, SE = 0.0125)

### MAGMA

To investigate the functional implications of the genetic variants associated with occupational status, we performed gene-based, and gene-set analyses using MAGMA.^47^ We used FUMA^92^ to annotate, prioritize, visualize, and interpret GWAS results, to run MAGMA on our summary statistics, and to map SNPs to genes. We tested whether the genes prioritized by FUMA were enriched for expression in 30 general tissue types (GTEx v8) and 53 specific tissue types (GTEx v8) using MAGMA’s gene-set analysis. We observed a strong expression in all brain tissues compared to other tissues. No other tissue showed significant enrichment for gene expression.

### MTAG

Multi-trait analysis of GWAS (MTAG)^48^ was used to meta-analyze all three occupational status measures with a secondary GWAS of household income in UKB and a secondary GWAS on educational attainment in UKB (for the validation subsample of siblings in UKB) or the third GWAS meta-analysis for education^6^ excluding 23andMe participants as well as the NCDS cohort (for the validation using the NCDS data). This allowed us to leverage the high genetic correlations between the occupational status measures and income/education (see above) to increase power and detect variants and improve prediction as outlined above and in SI 8.

### GSEM

We used the infrastructure provided by the GenomicSEM-package^42^ to compute LDSC-based genetic covariances and correlations between our occupational status measures and education and income. SNPs were included using similar criteria as specified for univariate LDSC. Covariance structures between the three measures of occupational status were used as input in a genomic structural equation model to analyze their loading on a joint factor of occupational status (see SI 9.1) We furthermore applied a multivariate genetic regression model to the genetic covariance matrix of each of our occupational status measures and cognitive performance, ADHD, openness to experience, risk tolerance, and neuroticism (see SI 9.3).

### Prediction analyses

Overall, we constructed three types of polygenic scores for each phenotype (see SI 8): (a) Pruning and thresholding using PRSice,^94^ (b) SBayesR,^95^ and (c) MTAG + SBayesR. In our prediction analyses, we residualized for sex, age (only in UKB), and 10 principal components before calculating the *R*^2^. For the within-family-analysis in UKBiobank we identified a sample of siblings and compute family-fixed effects regressions with both polygenic scores as well as phenotypes standardized beforehand and interpret the change in coefficients (see SI 10).

### Mediation analyses

NCDS respondents were asked at age 11 about the type of job they would like to do in the future. We coded these jobs to SOC2000, constructed their occupational status and ran mediation models in lavaan^96^ to quantify the share of the association between PGS, and occupational status that can be attributed to occupational aspirations. We tested a comprehensive multiple mediation model, introducing cognitive ability, internalizing behavior, scholastic motivation, and externalizing behavior as additional mediators (Supplementary Figure 8).

### Confounding analyses

Within NCDS, information on the paternal occupation at age 12 was used to estimate the correlation between paternal and offspring occupational status at various ages for all three measures. We combined the approach of scaling the variance explained by polygenic scores, outlined by Tucker-Drob (2017),^61^ and integrated it into a mediation model to test which share of the intergenerational correlation for each of the three metrics is confounded by the corresponding polygenic score if we assume that it only explains the amount of variance in our prediction analysis or the full SNP-heritability (r=0.14 for CAMSIS, r=0.1 for ISEI and SIOPS, see SI 12).

## Data availability

Genome-wide summary statistics will be publicly available upon publication on the GWAS Catalog website: https://www.ebi.ac.uk/gwas/downloads/summary-statistics. Access to the UKBiobank is available through the application with information available at: http://www.ukbiobank.ac.uk). Access to The National Child Development Study (NCDS) is available through the application as well at: https://cls.ucl.ac.uk/data-access-training/.

## Code availability

R-package ‘ukbjobs’ available at https://github.com/tobiaswolfram/ukbjobs. The package allows researchers to construct CAMSIS, ISEI, and SIOPS occupational scores directly from the UKBiobank data. No other custom code was used; all analyses and modeling were performed using standard software as described in the Methods section and in the Supplementary Information.

**Figure 8.**
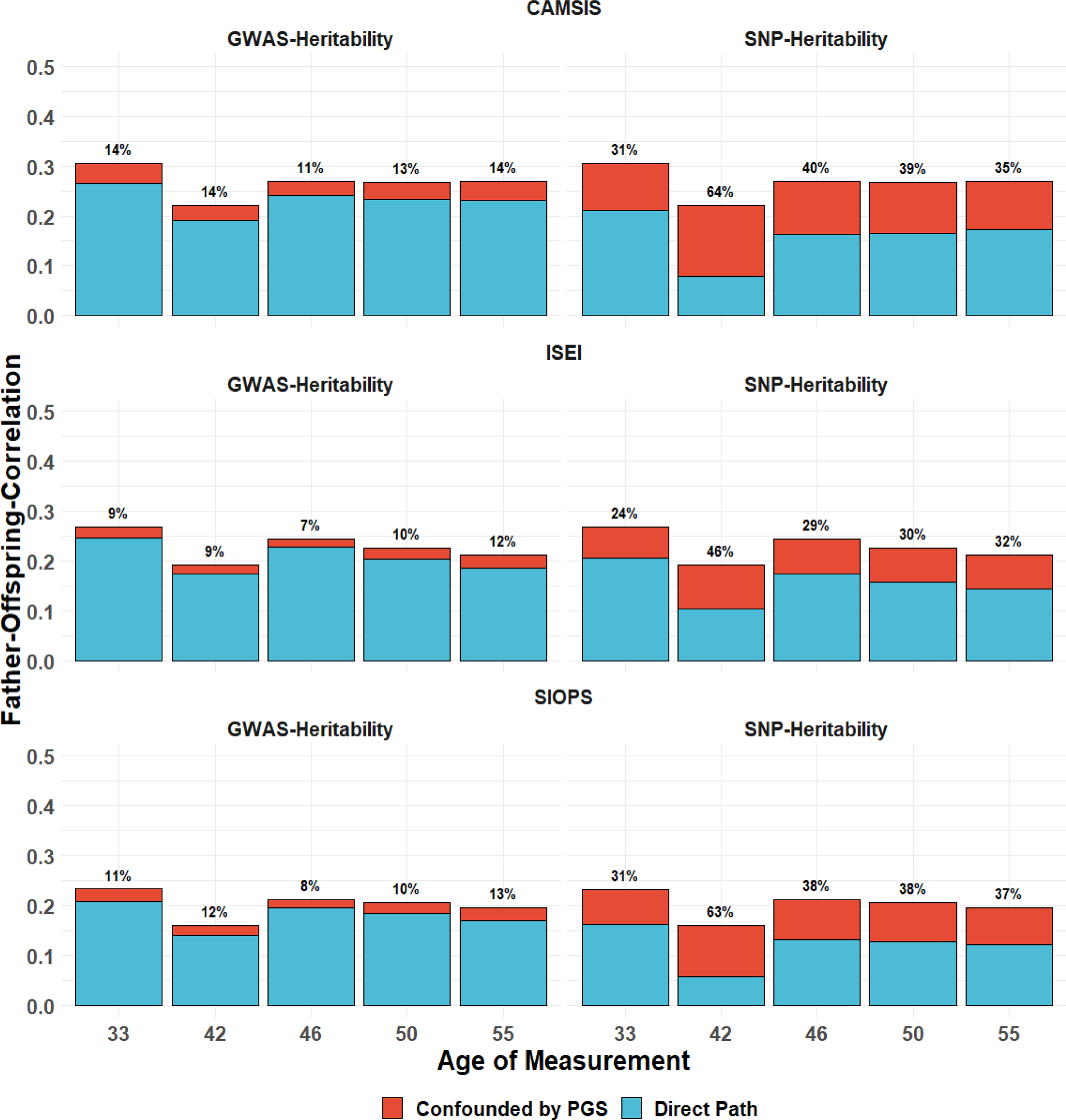
Depicting the percentage of genetic confounding in the intergenerational transmission of occupational status based on the predictive validity of polygenic scores (GWAS-Heritability) and an extrapolation of their effect to the variance explained by common SNPs (SNP-Heritability) in NCDS through the life course. N = 3,875; 3,835; 3,747; 3,550; 3,079 for CAMSIS at age 33, 42, 46, 50, 55 and 3,902; 3,797; 3,718; 3,522; 3,053 for SIOPS/ISEI.

## Acknowledgements

This research was conducted using the UK Biobank resource under application 32696 and NCDS under application GDAC_2021_16_TROPF. Funding for this project is from the European Research Council ERC Advanced Grant CHRONO (835079) & Leverhulme Trust (RC-2018-003) Leverhulme Centre for Demographic Science, to PI MCM. The study received ethical approval from the Department of Sociology, University of Oxford. The authors thank Dr. David M. Brazel for his important contribution and comments at the initial stage of the discovery.

## Author information

These authors contributed equally: Evelina T. Akimova, Tobias Wolfram

These authors jointly supervised this work: Felix C. Tropf, Melinda C. Mills

Contributions: M.C.M and F.C.T supervised the study. T.W. and F.C.T. wrote the paper, with extensive revisions and comments by M.C.M. and E.T.A. E.T.A and T.W. wrote Supplementary Information with comments by all authors. T.W. and E.T.A. conducted statistical analyses with input from X.D. on representativeness of occupations analysis. All authors reviewed and approved the final version of the paper.

Corresponding authors: Evelina T. Akimova, Tobias Wolfram

## Supplementary Information

**Supplementary Tables**

Supplementary Tables B1-B8.

