## Supplemental Information for "Polygenic predictions of occupational status GWAS elucidate genetic and environmental interplay for intergenerational status transmission, careers, and health"

#### Table of Contents


List of supplementary figures 3

List of supplementary tables 4

1. Frequently Asked Questions (FAQ) 5

2.Background 12

3.Measuring occupational status and prestige 13

3.Research plan 13

4.Phenotype definitions 14

5.Representativity of the UK Biobank compared with the Office of National Statistics (ONS) 14

6.Overview of GWAS analyses 16
- 6.1 Analyses 16
- 6.2 Sample inclusion criteria 16
- 6.3 Findings 17
- 6.4 Replication 17

7.SNP-heritability 18

8.Polygenic score calculation and prediction 18
- 8.1 Calculation of polygenic scores 18
- 8.2 Out-of-sample prediction 19

9.Uncovering genetic communality of occupational status and prestige with socio-economic and other measures using Genomic SEM 19
- 9.1 General factor of occupational status 19
- 9.2 General factor of socioeconomic status (SES) 21
- 9.3 Mediators between polygenic signals and occupational status 22

10.Direct and indirect effects 24
- 10.1 Parental Control design 24
- 10.2 Adoption design 25
- 10.3 Sibling design 25
- 10.4 The sibling design and assortative mating 25

11.Polygenic score prediction over the life course 26

12.Polygenic scores and the intergenerational transmission of occupational status 27

13.Polygenic score mediation by occupational aspirations and psychological traits 27

14.Polygenic scores and occupational status trajectories throughout the life course 29

15.Polygenic score associations with health outcomes 31
- 15.1 General health 31
- 15.1a Controlling for parental occupational status 32
- 15.1b Confounding of phenotypical effect of occupational status 33
- 15.2 Mental health 34
- 15.2a Controlling for parental occupational status 36
- 15.2b Confounding of phenotypical effect of occupational status and mental health 37

References 38

### List of supplementary figures

*Supplementary Figure 1. Distribution of latest occupational prestige measured by ISEI from both the UK Biobank sample and the ONS sample*

*Supplementary Figure 2. Distribution of latest occupational prestige measured by SIOPS from both the UK Biobank sample and the ONS sample*

*Supplementary Figure 3. Phenotypic correlation (upper right triangle) versus Genetic correlation (lower left triangle) of occupational prestige and status measures*

*Supplementary Figure 4. Path diagram of Confirmatory Factor Analysis (CFA) for a general factor of occupational status*

*Supplementary Figure 5. Phenotypic correlation (upper right triangle) versus Genetic correlation (lower left triangle) of occupational status measures and other SES indicators*

*Supplementary Figure 6. Path diagram of Confirmatory Factor Analysis (CFA) for a general factor of socioeconomic status (using CAMSIS as the occupational indicator)*

*Supplementary Figure 7. Incremental R-square of polygenic score predictions of occupational status over the life course, NCDS*

*Supplementary Figure 8. Mediation results of polygenic prediction of occupational status (controlling for parental SES), NCDS*

*Supplementary Figure 9. Genetic confounding of correlations from respondent's occupational status to general health*

*Supplementary Figure 10. Genetic confounding of correlations from respondent's occupational status to mental health*

### List of supplementary tables

*Supplementary Table 1. Results of multivariate genetic regression models - CAMSIS and potential mediators*

*Supplementary Table 2. Results of multivariate genetic regression models - SIOPS and potential mediators*

*Supplementary Table 3. Results of multivariate genetic regression models - ISEI and potential mediators*

*Supplementary Table 4. Effect size reduction when controlling for parental SES*

*Supplementary Table 5. Parameters of Best Fitting Growth Curve Models, NCDS*

*Supplementary Table 6. Parameters of Best Fitting Growth Curve Models (controlling for parental SES), NCDS*

*Supplementary Table 7. Associations between occupational status PGS and general health at various ages, controlling for sex and first 10 PCs*

*Supplementary Table 8. Associations between occupational status PGS and general health at various ages, controlling for father's occupational status at age 11, sex and first 10 PCs. Ratio denotes the ratio of the standardized beta-coefficient of the PGS without controlling for father's occupational status and the standardized beta-coefficient of the PGS when controlling for father's occupational status in the same sample of individuals for which paternal occupational status information was available.*

*Supplementary Table 9. Associations between occupational status PGS and general health at various ages, controlling for father's occupational status at age 11, sex and first 10 PCs*

*Supplementary Table 10. Associations between occupational status PGS and mental health at various ages, controlling for father's occupational status at age 11, sex and first 10 PCs. Ratio denotes the ratio of the standardized beta-coefficient of the PGS without controlling for father's occupational status and the standardized beta-coefficient of the PGS when controlling for father's occupational status in the same sample of individuals for which paternal occupational status information was available.*

### 1. Frequently Asked Questions (FAQ)

#### WHY THIS FAQ?

This study looks at occupation status and draws from multiple scientific approaches from the social sciences, molecular genetics, biostatistics and medical sciences. Given the controversial nature of examining differences in socioeconomic status and class in the context of the genome (see Box 1 main article), we wanted to create an accessible document for a broader audience to clarify what we are, and importantly, are not, doing with this study. It is aimed at those who are new to the scientific terminology and methods. Experts or those seeking more in-depth information and scientific references supporting our statements, should refer to our main article and the detailed Supplementary Material.

A **Glossary of terms** can be found at the end of this document.

#### WHAT DID WE STUDY?

The article by Akimova & Wolfram et al. (2023) examines one of the core topics of social stratification research, which is a field of research in the social sciences that studies and categorises groups of people based on socioeconomic factors like wealth, income, education, or occupation.

Moving beyond the previous focus on education, income, and wealth, we studied occupational status using three measures derived from decades of research in the disciplne of sociology, the: *International Socioeconomic Index (ISEI), Standard International Occupational Prestige Scale (SIOPS),* and *Cambridge Social Interaction and Stratification Scale (CAMSIS)*.

#### WHY STUDY THIS TOPIC NOW?

Socioeconomic status (SES) is a complex phenomenon influenced by behavior, biology, and the social environment. Understanding SES requires a multidisciplinary approach that identifies common drivers and addresses the multidimensional nature of their relationships with biology, health, social environments, and other behaviors.

Traditionally, biological and social processes of disease or complex behavior inheritance have been studied separately and therefore in a field-deterministic manner. However, the joint consideration of social and biological factors in a biosocial model, provides a more comprehensive scientific understanding. Advances in technology, such as genotyping and large datasets that incorporate genetic and environmental/behavioral information, have spurred a wide range of empirical inquiries, enabling improved modeling of complex behaviors and *traits*.

Social stratification is a central topic across the social and health sciences. Particularly, the intergenerational status transmission and reproduction within families has received great attention in the literature and previous research demonstrated that socioeconomic status is influenced by multiple factors including family, societal and historical contexts, and social norms. Next to the family environment, parenting behavior and other investments in children, genetic inheritance plays a role. For socioeconomic status measures such as educational attainment, nearly 4,000 *genetic variants* have been associated with the outcomes in previous studies. *Genetic variants* associated with income have also been established. Previous research has also shown that the third measure of occupational status, is not only driven by socio-environmental factors, but it also potentially has a biological and genetic basis. Twin studies suggest a *heritability of occupational status* of 0.30 to 0.40. The current study goes substantially beyond what we know about the genetics and biological factors associated with occupational status.

Though knowledge about potential genetic effects has led to various interpretations by social scientists, often in relation to a measure of merit. Thus, a quantitative exploration of factors through which social and biological predictors are linked enhances our understanding of the nature of the links between the genome and social stratification in general. This approach helps prevent potentially misleading interpretations of latent genetic measures in the context of questions regarding equality of opportunity.

To date, the majority of research on differences in socioeconomic outcomes has been studied using a socially determinist approach, focussing only the role of social and contextual factors on prediction. More recently, however, researchers have conducted *genome-wide association studies (GWASs)*, which scans the entire genome to discover the genetics related to socioeconomic and other complex behavioural outcomes. Previous *GWASs* were conducted for education and income (including household income). But someone might have high education, but not obtain a good job or high income. Or, someone might have low income and rise high in the occupational ranks. It is therefore interesting to understand whether there are genetic underpinnings of occupational status and how these genetics operate in relation to other socioeconomic outcomes, with health and across different environments, families and over time.

We therefore performed the first *GWAS* on measures of occupational status along with various follow-up statistical analyses in order to understand the nature of discovered correlations and the complex interplay between genes and environments.

#### WHAT WAS THE AIM OF THIS STUDY?

The aim of this study is to improve our understanding of occupational status attainment and transmission, specifically through the complex interplay between biological inheritance and social processes. To achieve this we:

1. Identified *genetic variants* associated with our measures of occupational status.
2. Utilized our genetic discovery results to create individual genetic scores (called polygenic scores), which we used to control for genetic associations when studying factors related to socioeconomic status, occupations, labor market, and occupational mobility.
3. Examined the genetic correlates of occupation in relation to other socioeconomic indicators,
4. Investigated the extent to which genetic associations of occupational status reflect the interplay between genetics, biology, family, social, and environmental factors.
5. Explored the potential mechanisms linking the genome and occupational status.

6. Comprehended the underlying structure of the discovered associations.
7. Explored the relationships between genes, occupational status, and (mental) health.
8. Introduce a life course perspective to examine how the genetic scores operate over time as individuals age and progress through their careers.

#### WHAT ARE THE BIG TAKE-AWAY FINDINGS?

In the **first genetic discovery study of occupational status** to date, we analyzed data from 273,157 individuals (130,952 males and 142,205 females) and i**dentified 106 *genetic variants*, including 8 newly associated** with the genetics of socioeconomic status.[1]

Our genetic predictor (*polygenic score*) **explains around 5-8% of the variability in occupational status** in the population, with analyses suggesting that *polygenic scores* based on larger samples may explain (depending on the measure) up to 11-15%. This is more than three times as much as found in a previous study that used a cruder measure of occupation.

The **intergenerational transmission of occupational status** only partly explain children's occupational status. Occupational status is only partly associated with *genetic variants*, and partly with the family background; the rest is neither family nor genes but unique circumstances and factors that we still need to measure. Notably, this is not specific to occupations, but to most complex disease, behavioral and social outcomes.

There is a **strong genetic interdependence among the three occupational status and a remarkably strong genetic correlation between other socioeconomic factors**, namely educational attainment and income. It is noteworthy that the genetic correlations observed among these socioeconomic indicators exceed the phenotypic correlations by a factor of two to three. Such a pattern is highly unusual in genomics and not observed for behavioral phenotypes or diseases.

**Cognitive skills, scholastic motivation, occupational aspiration, personality traits, and behavioral disinhibition (represented by ADHD) play a significant role** in driving the association between genetics and occupational status.

**Genetic predictions can vary throughout an individual's life course**, potentially influencing their labor career trajectory, suggesting that future behavioural and disease-related research should examine how the association of genetics vary as individuals age.

**Social environmental factors, such as the family that someone grows up in, has a strong impact** via what is called gene-enviornment correlations. **When parental occupational status and data from adoptees is examined, the polygenic score's impact diminished by roughly 25%.** A greater reduction of over 50% was observed when predicting differences in occupational status

[1] It is important to emphasize that there is no single definitive gene responsible for the complex outcome of occupational status, nor are there direct causal genes that determine an individual's occupational status. Our intention was not to identify such genes, as occupational status is a result that is influenced by multiple factors beyond direct genomic influence and functional biology. However, we consider it crucial to underscore this concept to ensure that our findings are interpreted accurately and responsibly.

between siblings in the same family. This disparity may stem from the strong socioeconomic status-based assortative mating over generations that can bias within-family estimates downwards. Thus, our findings suggest that **gene-environment correlations likely account for approximately 25% of the discovered estimates.**

#### HOW DOES THE CURRENT STUDY EXTEND OUR KNOWLEDGE?

The current study contributes to our understanding in several significant ways, namely:

1. **First and largest genetic discovery for occupational status** to date including 273,157 individuals (130,952 males, 142,205 females) where we used sociologically theorized measures of occupational status.
2. We **include an X-Chromosome analysis**, allowing us to uncover one additional novel locus.
3. **We conduct various novel quantitative downstream analyses**, including:
   A) analyses on genetic confounding of the intergenerational transmission of occupational status,
   B) identifying mechanisms linking the genome and occupational status, and
   C) investigating genetic associations with occupational status across the life course.
4. **Embrace a comprehensive and interdisciplinary perspective** in the study of social stratification, mobility, and status transmission. By delving into the underlying latent factors of individual socioeconomic status indicators, we can cultivate a deeper understanding of socioeconomic status and its wider societal implications.
5. **Demonstrate it is crucial to comprehend the role of *indirect effects* and *passive gene-environment correlations*** in unraveling this complex puzzle. Both of these aspects are pointing towards the importance of the interplay between genetic variants and (social) environments and its complexity.
6. **Understanding the causes and consequences of *assortative mating*** on these relationships is of utmost importance. By exploring these aspects, we can gain valuable insights into the intricate dynamics that shape social stratification and contribute to a more holistic understanding of socioeconomic status.

#### HOW DID WE STUDY IT?

The primary analysis we conducted is called a *Genome-Wide Association Study or GWAS* (pronounced gee-was), which is a search across the entire human genome, examining each genetic locus (or region) one by one to see if there is a relationship (or what we call an association) between our outcomes and a particular *genetic variant*. *Variants* refer to a specific region of the genome, which differs between two genomes. Different versions of the same *variants* are termed alleles and a *SNP* (pronounced SNiP; *single-nucleotide polymorphism*) can have two alternative bases or alleles (C and T).

In other words, we study DNA *variants* that distinguish us from each other. Humans are 99.9% identical to each other, and it is the 0.1% by which we differ that makes us all genetically unique. A small subset of the 0.1% by which we differ genetically is anticipated to influence reproductive behavior.

Moreover, a comprehensive interdisciplinary study such as this one demanded multiple analytical approaches, we:

- investigated the functional implications of *genetic variants* associated with occupational status through gene-based and gene-set analyses using MAGMA technique.
- employed multi-trait analysis (MTAG) to meta-analyze occupational status measures with household income and educational attainment.
- utilized genomic structural equation models (GSEM) to analyze the joint factor of occupational status, cognitive performance, ADHD, openness to experience, risk tolerance, and neuroticism.
- *polygenic score (PGS)* construction and prediction involved producing various scores, testing out-of-sample prediction, and assessing *population stratification* using LD score regression.
- applied sibling and adoption models to disentangle direct, indirect, and demographic effects.
- conducted mediation and confounding analyses.

#### WHO ARE WE AND WHO FUNDED THIS STUDY?

We are grateful for the contribution of all UK Biobank and NCDS participants to this scientific study. The UK Biobank was approved under application 32696 and the NCDS data use was approved under application GDAC_2021_16_TROPF. Rsearch funding support was provided by the ERC (European Research Council) Advanced Grant to Melinda Mills in 2019 for the CHRONO project (number 835079) and the Leverhulme Trust, Leverhulme Center for Demographic Science (LCDS, grant number N/A).

#### ADDITIONAL GENERAL QUESTIONS

##### Are the genetic associations small or large?

Occupational status, similar to other socioeconomic measures, is a complex outome that is not only genetically based but shows a complex interplay with individual and socio-environmental contextual factors. Genetics is only one piece of this larger puzzle and in this study we only examine one type of *genetic variants* (*SNPs*) and consider only one of the many possible biological and genetic ways in which individuals may vary. This does not impact the importance of the findings, since one single factor or variable ever fully explains complex outcomes.

##### Could genetic results alone be used at the individual level to predict someone's occupational status?

With the exception of some diseases, to date, genetic scores alone are usually not useful to predict complex individual disease and behavioural outcomes. When we are examing complex behavioural outcomes each individual *SNP* or *genetic variant* has a small effect, so prediction of using genetic results alone is not possible. Even if we combine the information contained in the more than 10 millions *genetic variants* that we studied together into a genetic predictor, we predict 5-8% of the variance across individuals. With larger samples we see that the ceiling of prediction is likely more in the range of 11-15% (depending on the measure of occupational status used). Extrapolating findings from other *traits*, more granular and detailed genetic data (on structural variation, insertions, deletions and rare *variants*) might further increase this ceiling.

However, even the 'gold standard' social science predictors when entered alone as a single variable in a regression equation also have low predictive power, generally under 10%. It is therefore unhelpfully reductive to only enter one single variable as a predictor without considering additional factors.

In reality, complex outcomes are a culmination of multiple factors such as genetics, parental background, lifestyle, level of education and national institutional configurations that constrain or enable behavior. We have in the past shown that the explanation of genetics can vary across country and time.

##### Is it nature or nurture?

That is a false dichotonomy and it is neither nature or nature but rather nature *and* nurture. Occupational status – similar to other socioeconomic status measures - is not nature versus nurture, but rather a combination of both. Just as complex diseases such as obesity or diabetes are neither purely genetically or socially determined, occupational status relates not only to biological factors, for example, influencing abilities or behavior, but also have a strong social and environmental component in that they are driven by partners, and simultaneously shaped by the social, cultural, economic and historical environment. Genetic factors partly influence the first two factors of biological ability and behavior, complemented by social and environmental influences which also filter the types of behavior that are possible in the historical environment (e.g., via legislation, labour market structure, social norms).

##### Are there societal or medical implications of this study?

Societal, maybe, but medical, extremely unlikely. In the longer term, this study offers a better understanding of the genetic architecture and responsible observable *traits* for occupational status. It equips social scientists to take genetic effects into account in their study and reduce bias due to genetic effects in their study of interest. But it also alerts medical and health researchers that genetic scores for complex *phenotypes* are also picking up considerable social environmental and family effects and that *polygenic score* prediction vary by age.

We reveal changing genetic effects across the occupational career which opens an interesting puzzle for life course researchers with the potential for discoveries of lifestyle factors interacting with genes. Our analyses of the relationship between occupational status, genes and, (mental) health have ramifications for public health as we can demonstrate that ignoring one of the dimensions produces a biased view on the other one. Furthermore, it is important to understand whether and which proportion of these *traits* are driven by genetic, behavioral and environmental factors. The fact that we also found evidence that genetic influences are much more shared than it's observed for different status measures suggests that continued research in this area is warranted to aid a better understanding of what makes the difference between income, education, and occupational status.

##### What are the limitations of this study?

Although we open up new avenues of research, there are limitations and which are not exhaustive or exclusive to this type of *GWAS* study, the central ones are listed here:

- Sample sizes are smaller than for example in the study of educational attainment which makes a direct comparison of results harder.
- Focus on European-ancestry individuals only, a problem we have highlighted elsewhere.

After conducting a scientometric review of all *GWAS* and realizing that 72% of genetic discoveries come from 3 countries, we set up the GWASDiversityMonitor described in our Nature Genetics article. We have also considered problematic biological race and genetic essentialism narratives.

Data availability
The summary statistics will be available on the GWAS Catalog website. The UKBiobank is available through application with information available at: http://www.ukbiobank.ac.uk/.

#### GLOSSARY

From Mills et al. (2020) An Introduction to Statistical Genetic Data Analysis, Cambridge: MIT Press.

**Phenotype or trait.** The observable characteristic of an individual, ranging from physical traits (hair colour, height) to disease status (diabetic) to behaviour (risk-taker, age at first sexual intercourse, educational attainment).
**Genotype.** Describes part of an individual's DNA that influences their phenotype.

**Genome-wide association study (GWAS).** A GWAS is designed to adopt an unbiased hypothesis-free approach to discover genetic variants are associated with a trait. They often combine data from multiple studies to gather the largest sample possible. An updated and searchable list of all GWAS discoveries to date can be found at www.gwasdiversity.com, linked to this article, with summary statistics available at the GWAS Catalog.

**Single-nucleotide polymorphism (SNP).** A common variation in a single nucleotide (i.e., A, C, G, or T) that occurs at a specific position in the genome. A SNP exists as two different forms (e.g., A vs. T). These different forms are called alleles. A SNP with two alleles has three different genotypes (e.g., AA, AT, and TT).

**Genetic variant.** Refers to a specific region of the genome that differs between two genomes.

**Heritability.** A population measure defining the proportion of variance in a phenotype explained by genetic variance within a population. We can differentiate between broad-sense heritability, including both additive and non-additive genetic effects such as epistasis and dominance, and narrow-sense heritability focusing on additive genetic effects only.

**SNP-heritability.** The fraction of phenotypic variance of a trait explained by all SNPs in the analysis. Usually less than the narrow-sense heritability as it does not take rare variants and structural variation into account.

**GWAS-heritability.** The fraction of phenotypic variance of a trait explained by genome-wide significant genetic variants—sometimes also by polygenic scores based on GWAS findings.

**Assortative mating.** In genetic research refers to a mating structure in which pairs of individuals that are (genetically) similar to each other mate with a higher probability than expected under random mating. Assortitave mating is an important concept for statistical genetics; it biases heritability estimates.

**Polygenic score (PGS).** A single quantitative variable that summarizes genetic association to a phenotype by combining multiple genetic variants and their associated weights, derived from a GWAS.

**Indirect genetic effects.** Refers to situations when environmental influences which are important for complex outcomes and phenotypes are also associated with individual's genotype. In such instances, enviroments referred as 'mediators' of the link between genetic variants and traits.

**Gene-environment correlations.** This is the process by which an individual's genotype influences or is associated with exposure to the environment.

**Gene-environment correlations.** Defines an interplay between a gene and an environmental factor in which the effect of the gene on a phenotype is modifiable by the environment, and vice versa.

**Population stratification.** The presence of multiple subpopulations (e.g., individuals with different ancestral background) in a study. Because allele frequencies can differ between subpopulations, population stratification can lead to false positive associations and/or mask true associations. An example is the chopstick gene, where a SNP, due to population stratification, would be wrongly assumed to be a true association due to differences in allele frequencies of those of Asian and European ancestry who have a different usage of chopsticks for purely cultural rather than biological reasons.

### 2. Background

Occupational status – including its prestige and other sociological factors - is a crucial component of socioeconomic status (SES) and correlates with many physical and mental illnesses as well as longevity.[1,2] Explanations for the SES-health gradient have also been examined in the context of genetics. Recent Genome-Wide Association Studies (GWASs) have identified the genetic contributions to economic status for education[3-5] and income.[6,7] Research on the genetic basis of occupation has to date focused mainly on entrepreneurship[8] and occupation-related stress and exhaustion.[9,10] Little is known about the genetic associations with occupational status and prestige, which is related to an individual's education and income level, and signals their social standing.[11]

The current study is the first GWAS of expert measures of occupation, extending previous work in several ways. Our analyses focus on the International Socioeconomic Index (ISEI), the Standard International Occupational Prestige Scale (SIOPS), and the Cambridge Social

Interaction and Stratification Scale (CAMSIS). In the following we detail our analyses which had been preregistered February 2nd, 2021 (https://osf.io/djbr2/) and updated for replication (https://osf.io/x6va5).

### 3. Measuring occupational status and prestige

The three approaches from the sociological tradition to measuring occupational status and prestige either consider socioeconomic differences between occupations, interoccupational social interaction or ascribed prestige of different jobs.[12]

Dating back to their earliest inception in the 1950s[13,14] *socioeconomic difference*-based indices measure the "attributes of occupations that convert a person's main resource (education) into a person's main reward (income)".[15] The most used measure of this tradition is the *International Socioeconomic Index* (ISEI[15]), which is a measure of prestige or status constructed from scaling weights that maximize the (indirect) influence of education on income through occupation.

More focussed *prestige*-based measures on the other hand are simply the result of public opinion surveys in which representative samples of the population are tasked with ranking occupations by their relative social standing, emerging at a similar time as socioeconomic difference-based indices (e.g., Nakao and Treas 1992[16]). Treiman (1977)[17] showed in an extensive analysis that prestige-based measures were surprisingly constant over time and cultures, cementing their use in social scientific research. His *Standard International Occupational Prestige Scale* (SIOPS or Treiman-prestige[18]) remains one of the most commonly used metric in this tradition.

Lastly, occupational status indicators derived from *social interaction* focus on the heterogeneity of associations between occupants of different jobs, following the tradition of Warner, Meeker, and Eells (1949)[19] and Laumann and Guttman (1966).[20] They are based on the idea that differential association is a function of social stratification, as members of a group are more likely to interact with members of that group than with members of other groups. Thus, acquaintances, friends, and spouses are much more likely to be selected from within the same group than an outgroup. A group of Cambridge sociologists reversed this approach to measure occupational structure based on interactions. The *Cambridge Social Interaction and Stratification Scale* (CAMSIS) measures the distance between occupations based on the frequency of social interactions (operationalized as husband-and-wife combinations) between them.[21]

### 3. Research plan

Analyses followed the pre-registration uploaded by Brazel, Ding and Mills to the open science framework on February 2nd, 2021 and updated in February 2023 also including CAMSIS (https://osf.io/x6va5).

The focus was on the most recent occupation the participant held. We studied ISEI, SIOPS, CAMSIS, the (within-family) prediction of polygenic signals and beyond. Analysts (ETA, TW) achieved genetic correlations of 1 in their exploration.

### 4. Phenotype definitions

Discovery is conducted in the UK Biobank, which is a large prospective cohort study in the United Kingdom (UK), following over 500,000 volunteers who were between 40 and 69 years of age at the time of their recruitment between 2006 and 2010.[22,23] The definition used in each research centre conducting the analysis is now outlined below.

Current or most recent occupation is treated as a continuous measure. UK Biobank respondents were asked to provide job titles for the current or the most recent job held. The job information was coded using the four-digit UK Standard Occupational Code version 2000 (SOC2000). We built a procedure to link the UK SOC2000 to ISCO-88(COM), and then derive ISEI and SIOPS from ISCO-88(COM). All phenotypes are inverse-normal rank transformed before analysis. The CAMSIS-based status could be directly merged using the data available from Lambert and Prandy (2018).[24] ISEI and SIOPS (as provided by the R-package "strat"[25]), however, use the less granular ISCO-88[26] scale, so a mapping from ISCO to SOC2000[27] was employed. If multiple job codes for a respondent were available, the newest was used.

For some of our analyses, we rely on the genotyped subsample of the National Child Development Study (NCDS). Here, the SOC2000 code of the respondent's occupation (as well as their father's when they were 11 years old), is available as well, so a similar procedure is applied.

We also measured other SES dimensions such as income as done by Hill et al. (2019)[7] using a coarse, 5-level ordinal household income variable. Educational attainment was defined as years of education and coded according to the scheme provided by Lee et al. (2018).[3]

### 5. Representativity of the UK Biobank compared with the Office of National Statistics (ONS)

Supplementary Figure 1 and Supplementary Figure 2 show the distribution of latest occupational status measured by ISEI and SIOPS from both the UK Biobank sample and the Office of National Statistics (ONS) sample. The purpose of comparing the distribution of the two samples is to check whether our phenotypic distribution of occupational status is broadly representative of the national population. It's categorised into eight (ISEI) and seven (SIOPS) groups.

We also compare the sex ratio of each occupational group in the two samples. Supplementary Figure 1 and Supplementary Figure 2 demonstrate that the UK Biobank sample is largely in line with the general population. However, we found that the UK Biobank sample in general under-represents individuals in the lower end of the occupational prestige and over-represents those in the higher end of the occupational prestige. In addition to potential sample selectivity of the UK Biobank, this is also attributed to the fact that the UK Biobank sample covers individuals over 40 years old whereas the ONS data source samples the entire labour force. The sex ratio of the UK Biobank and UK population sample are highly correlated.

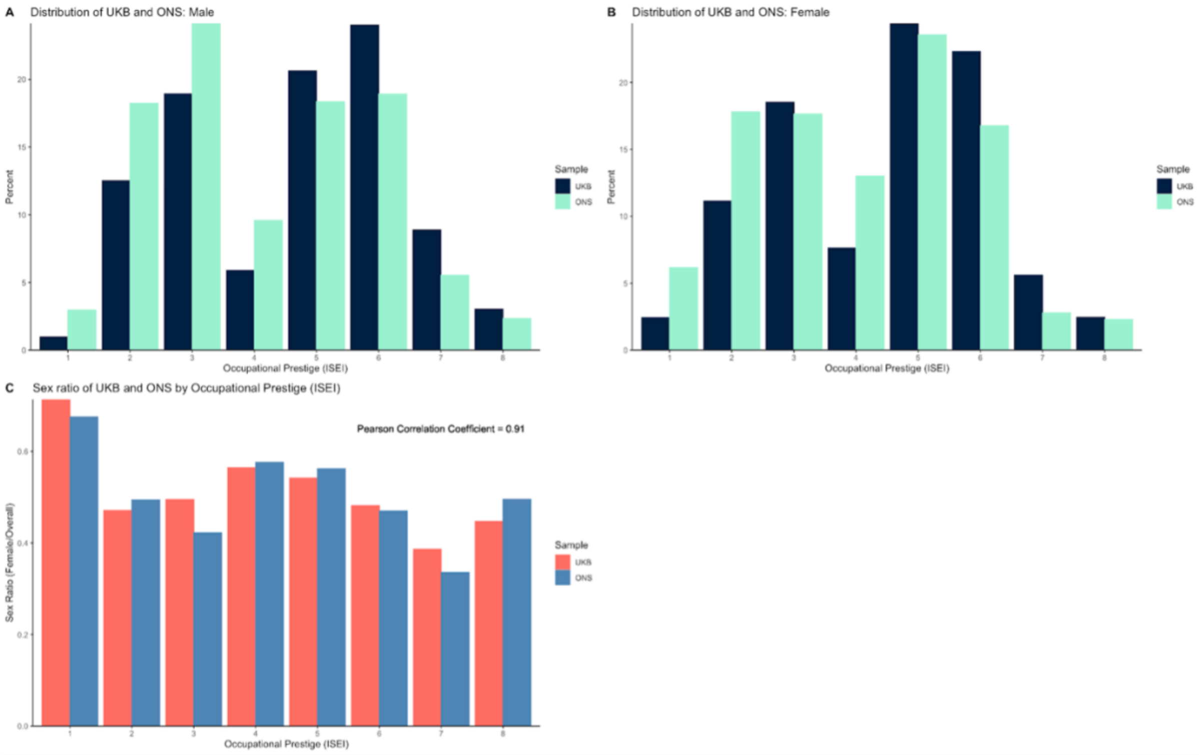


*Supplementary Figure 1. Distribution of latest occupational prestige measured by ISEI from both the UK Biobank sample and the ONS sample*

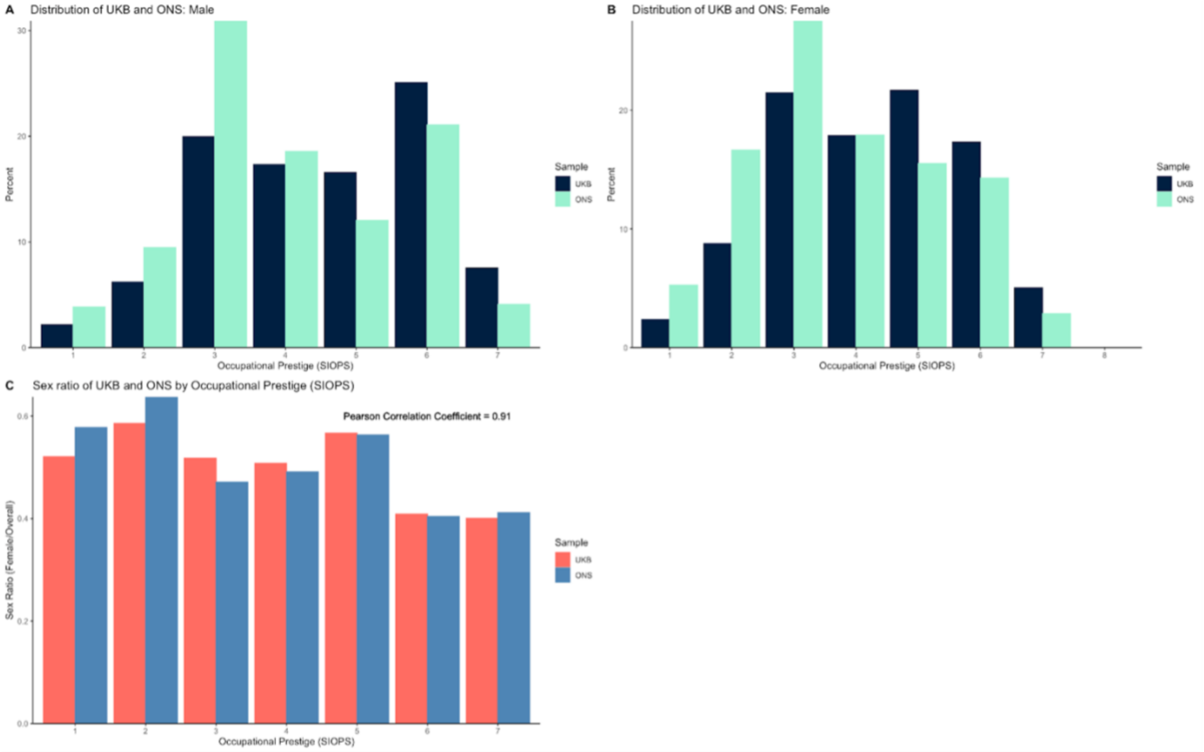


*Supplementary Figure 2. Distribution of latest occupational prestige measured by SIOPS from both the UK Biobank sample and the ONS sample*

### 6. Overview of GWAS analyses

For discovery, we followed the analysis plan first uploaded by Brazel, Ding, and Mills to the open science framework on February 2nd, 2021, which was updated including CAMSIS (https://osf.io/djbr2/) in February 2023 (https://osf.io/x6va5).

#### 6.1 Analyses

All calculations are based on mixed model association tests as implemented in the program FastGWA,[28] with association testing based on v3 imputed data. Following the pre-posted open science analysis plan in each regression, the following covariates are included: The first 10 genomic principal components, age at assessment and age$^2$, UK Biobank assessment centre at recruitment, sex, genotyping array (BiLEVE or Axiom) on the sample of European ancestry. Chromosomes are analysed separately. To speed up the calculation of summary statistics, a minimum MAF filter of 0.01 was imposed, leaving 10.2 million SNPs for the analysis.

#### 6.2 Sample inclusion criteria

Individuals were included from "genetically white British Ancestry" who had information for the phenotype in question, all relevant covariates, no mismatch between submitted and inferred gender, no outlier for heterozygosity and missingness, no evidence for sex chromosome aneuploidy and if they were present in all relevant genetic datasets (unimputed, imputed autosomal and imputed X chromosome), leaving in total 273,157 (130,952 males, 142,205 females) and 271,769, (130129 males, 141640 females) individuals for the occupational status phenotypes (CAMSIS and SIOPS/ISEI) and 353,673 (1692,01 males, 184,472 females) and 404,420 (185,632 males, 218,788 females) individuals for the secondary analyses (household income and education), respectively. For all analyses based on the sibling sample of the UK Biobank, separate GWAS using the same specifications with the respective individuals removed were conducted.

#### 6.3 Findings

After inflating the standard errors by the square root of their respective intercepts from LD Score regressions, our GWASs identified 106 genetic variants for CAMSIS including 56 found also for ISEI and 51 for SIOPS based on an $R^2$-threshold of 0.1 and a window-size of 1000kb, one of which (only significant for CAMSIS) was found on the X-chromosome (see Figure 2 in the main text for the Manhattan plot of the autosome). In exploratory sex-specific GWAS, no separate hits emerged, with genetic correlations being very close to and not significantly different from one.

#### 6.4 Replication

To validate our findings, we replicated our top hits using the genotyped subsample of the National Child Development Study (NCDS), an ongoing British Birth Cohort Study which started in 1958 and includes ~6500 individuals with both genetic and phenotypical information.

We replicate our findings on CAMSIS, which includes the discovery for ISEI and SIOPS, exhibited the highest SNP-heritability (see SI 7) and achieved the best polygenic prediction (SI 8), showing a genetic correlation not significantly different from 1 to ISEI and SIOPS (SI 9). To keep in line with the age range of the discovery sample, we used occupational information later in life (age 50). After filtering for standard quality control measures like those in the discovery, the final dataset included 4,899 individuals with 10 PCs and sex as covariates.

Restricting our pre-clumped summary statistics to SNPs found in the NCDS, we were able to match 103 of the 105 autosomal independent genome-wide significant variants (or variants in LD to them) between both datasets for replication.

Considering the limitations imposed by the smaller sample size of the NCDS dataset, we turned to the methods presented by Okbay, Beauchamp, Fontana, Lee, Pers, Rietveld et al. (2016, SI 1.8.3),[29] to assess the expected sign concordance and significance of our results, allowing us to estimate the expected performance of our replication while considering the standard errors in both discovery and replication datasets.

The probability that the SNPs have the same sign in both discovery and replication datasets, $P(\text{match})$, is :

$$P(\text{match}) = \Phi\left(\frac{-|\beta|}{\sigma_{\text{GWAS}}}\right)\Phi\left(\frac{-|\beta|}{\sigma_{\text{replication}}}\right) + \left(1 - \Phi\left(\frac{-|\beta|}{\sigma_{\text{GWAS}}}\right)\right)\left(1 - \Phi\left(\frac{-|\beta|}{\sigma_{\text{replication}}}\right)\right)$$

where $\beta$ is the vector of winners curse corrected effect sizes froms the discovery, $\sigma_{\text{GWAS}}$ the vector of associated standard errors in the discovery and $\sigma_{\text{replication}}$ the vector of standard errors in the replication. Then $\Sigma P(\text{match})$ is the expected number sign-concordant SNPs.

The NCDS replication significantly outperformed expectations under the null hypothesis that discovered signals are not robust. At $p < 5 \times 10^{-8}$, the expected sign concordance was 54.2 under the null while the actual match was 67. At the suggestive significance level of $p < 5 \times 10^{-6}$, the expected sign concordance was 199.2, and the actual match was 255.

The probability of significant hits on a given $\alpha$-level, $P(\text{sig})$, can be computed as:

$$P(\text{sig}) = \Phi\left(\frac{-|\beta|}{\sigma_{\text{replication}}} + \Phi^{-1}\left(\frac{\alpha}{2}\right)\right) + \left(1 - \Phi\left(\frac{-|\beta|}{\sigma_{\text{replication}}} - \Phi^{-1}\left(\frac{\alpha}{2}\right)\right)\right)$$

Again, we can sum over the vector $\Sigma P(\text{sig})$ to get the expected number of hits.

Demonstrating the consistency of our findings, the number of significant hits at $\alpha$ = 0.05 was in line with expectations: At $p < 5 \times 10^{-8}$, the expected match was 5.2, and the actual match was 5 and at $p < 5 \times 10^{-6}$, an actual match of 24 was achieved, compared to an expectation of 19.4.

### 7. SNP-heritability

SNP-heritability for all three primary (ISEI, SIOPS, CAMSIS) and the two related SES phenotypes (education, household income - analysed in the UK Biobank) were computed using LDSC[30] from GWAS summary statistics. All measures exhibit SNP-heritability significantly larger than zero, as shown in Supplementary Figure 3 and Figure 3 in the main text. The values for household income and educational attainment are comparable to SNP-heritability previously published in the literature.[3,7] Heritability of ISEI and SIOPS is 0.10 to 0.11 and larger in CAMSIS, comparable to that of educational attainment, with 0.146.

### 8. Polygenic score calculation and prediction

#### 8.1 Calculation of polygenic scores

Overall, we calculated three types of polygenic scores for each phenotype:

1. **Pruning and thresholding polygenic scores using PRSice.**[31]
   Polygenic scores were calculated using a prespecified threshold of p = 0.5 in the sample and with the software default values for clumping (250kb window; $r^2$=.1).
2. **SBayesR[32] polygenic scores.** Polygenic scores were calculated using the software default values (pi = 0.95,0.02,0.02,0.01, gamma = 0,0.01,0.1,1) and a shrunk sparse LD-matrix computed based on 1.1 million common SNPs in a random sample of 50K unrelated individuals of European ancestry in UK Biobank provided at https://zenodo.org/record/3350914#.XyFfnC17G8o.[32]
3. **MTAG+ SBayesR polygenic scores.** We calculated this type of polygenic scores using the same specifications as in SBayesR polygenic scores but instead it is based on MTAG results[33] from GWAS on all occupation scores and the secondary GWAS on income and education. To maximize our predictive power, in the NCDS the EA3 GWAS[3] (excluding NCDS) is used instead of the secondary UKB education GWAS.

#### 8.2 Out-of-sample prediction

For all phenotypes, we use OLS regression models and report the incremental R-squared over a baseline model consisting of 10 principal components, sex and age. Accordingly, we investigated the predictivity of polygenic scores in the subsample of siblings in the UK Biobank (N = 24,579 for CAMSIS and 24,472 for ISEI and SIOPS, we ran additional GWAS excluding these observations from the sample) and in the genotyped subsample of the National Child Development Study (NCDS), an ongoing British Birth Cohort Study started in 1958. For the NCDS, observations were pooled over all waves starting at age 33 (N = 5,389; 5,312; 5,211; 4,902; 4,263 for CAMSIS at age 33, 42, 46, 50, and 55, N = 5,449; 5,293; 5,197; 4,892; 4,252 for ISEI/SIOPS). Figure 4 in the main text demonstrates the results of performance of different polygenic scores.

### 9. Uncovering genetic communality of occupational status and prestige with socio-economic and other measures using Genomic SEM

#### 9.1 General factor of occupational status

We observed strong correlations between all three occupational prestige and status measures. Supplementary Figure 3 below illustrates an extremely high genetic correlation between CAMSIS, ISEI, and SIOPS (lower triangular), much stronger than implied by their phenotypic correlations (upper triangular).

Using genomic structural equation modelling (R package GenomicSEM)[34] on the genetic correlation matrix, we find clear evidence for a general factor of occupational status (Supplementary Figure 4) with extremely high loadings of all three measures. It is much higher than a phenotypic exploratory factor analysis in the same sample (path coefficients of 0.86, 0.96 and 0.94, for CAMSIS, ISEI and SIOPS, respectively, not shown).

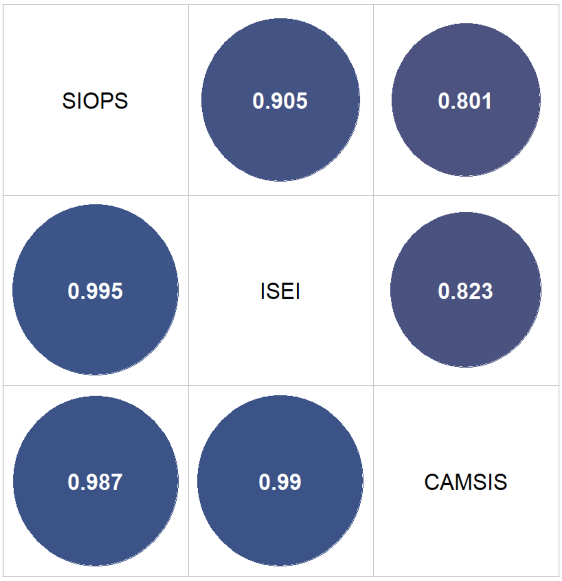


*Supplementary Figure 3. Phenotypic correlation (upper right triangle) versus Genetic correlation (lower left triangle) of occupational prestige and status measures*

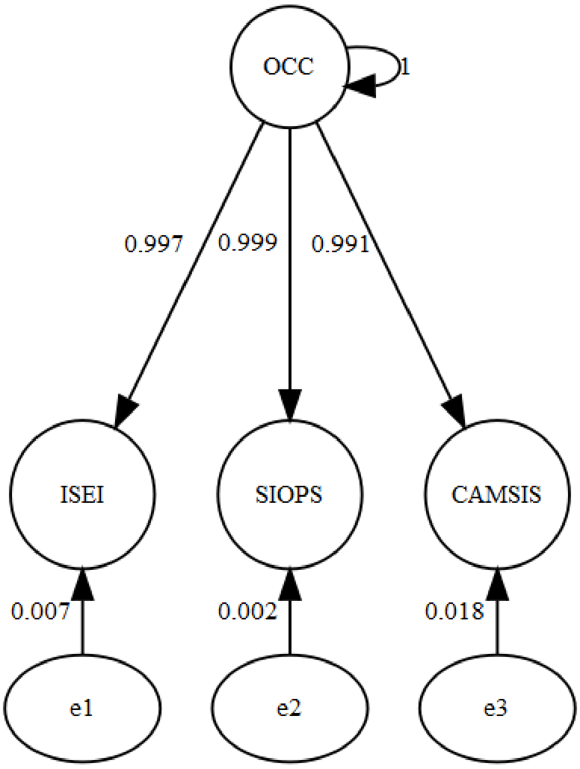


*Supplementary Figure 4. Path diagram of Confirmatory Factor Analysis (CFA) for a general factor of occupational status*

#### 9.2 General factor of socioeconomic status (SES)

In addition to exhibiting strong genetic commonality, Supplementary Figure 5 shows that genetic correlations of CAMSIS, ISEI and SIOPS with educational attainment and household income (lower triangle) are without exception in the extreme as well. This is in particular evident in contrast to the phenotypic correlations (upper triangle), which are existent and positive, but much smaller.

This implies the existence of a common factor of socioeconomic status, consisting of occupation, education, and income. This is again supported by factor analysis using genomic SEM (Supplementary Figure 6).

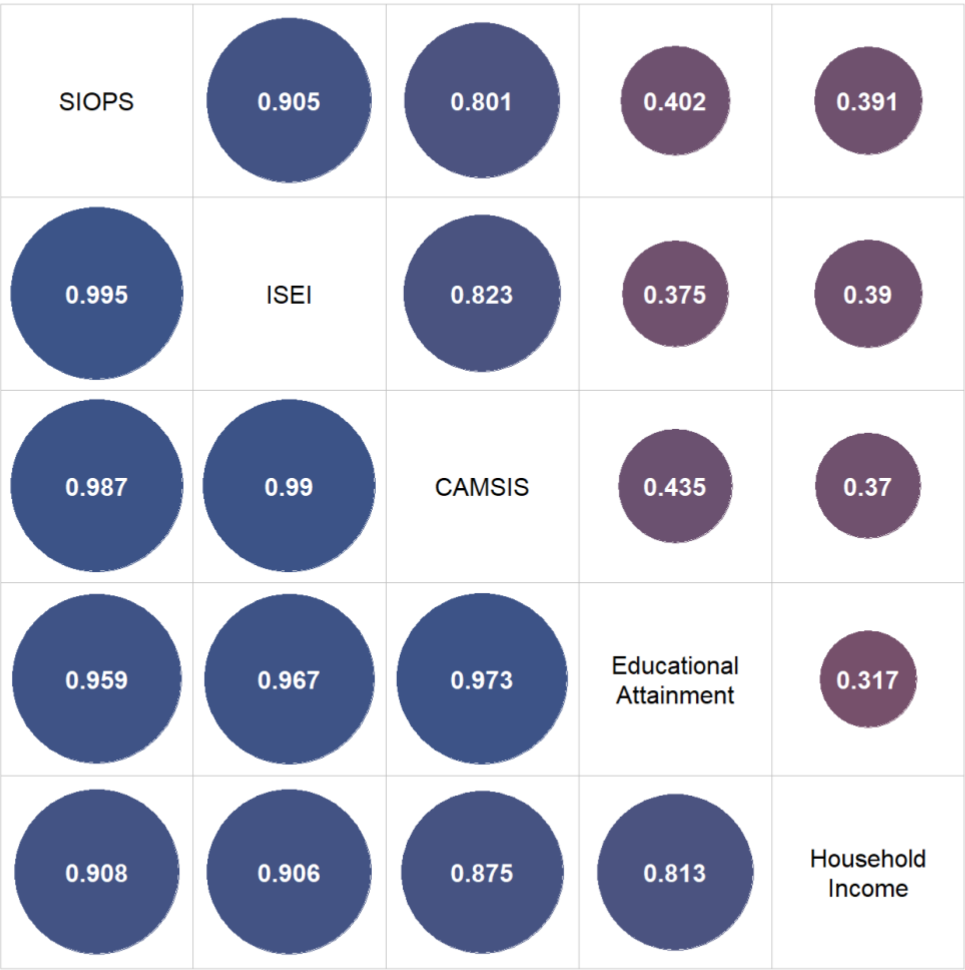


*Supplementary Figure 5. Phenotypic correlation (upper right triangle) versus Genetic correlation (lower left triangle) of occupational status measures and other SES indicators*

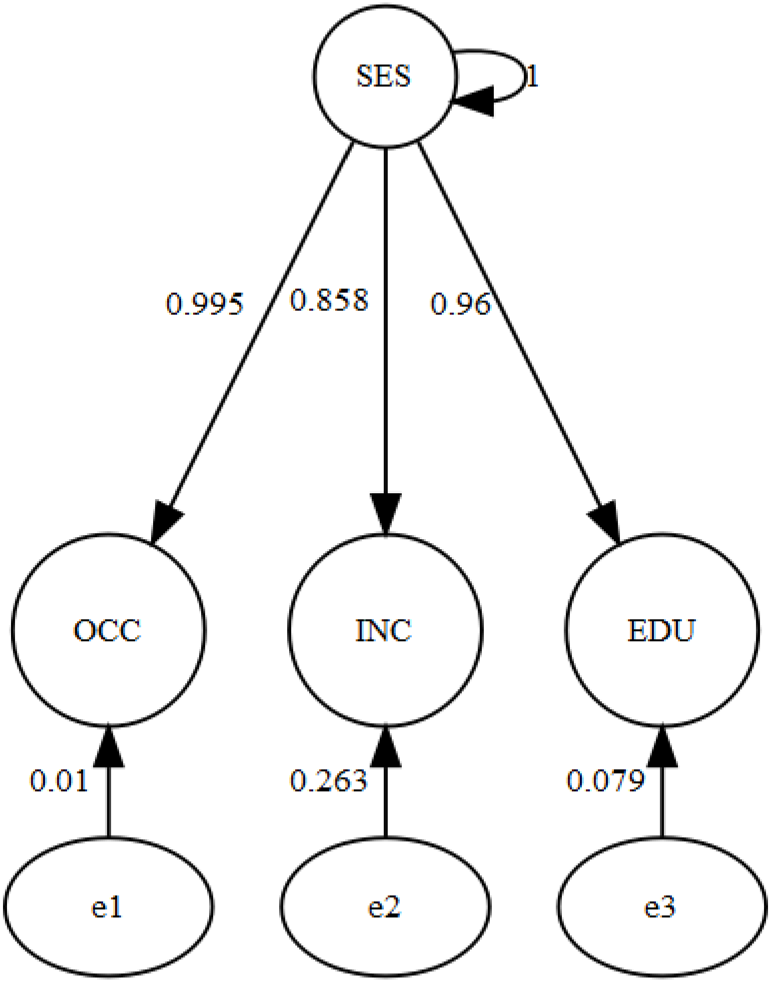


*Supplementary Figure 6. Path diagram of Confirmatory Factor Analysis (CFA) for a general factor of socioeconomic status (using CAMSIS as the occupational indicator)*

#### 9.3 Mediators between polygenic signals and occupational status

What biologically proximal traits are responsible for the common factor underlying genetic variation in socioeconomic status? Evidence from twin studies points to the direction of cognitive and noncognitive traits acting as mediators,[35] showing for example that the heritability of education is mostly due to a variety of noncognitive factors and intelligence,[36] all of which show substantial genetic influence.[37-40]

We validate these findings by identifying five causally upstream traits from the genetic literature that we expect to mediate the general genetic factor of socioeconomic status, namely cognitive performance,[3] ADHD (as a proxy for behavioural disinhibition),[41] openness to experience,[42] risk tolerance,[43] and neuroticism[44] and fitting a multivariate genetic regression model[45] in Genomic SEM.

Results presented in Supplementary Table 1 - Supplementary Table 3 show the respective regression results, which indicate that overall, almost 70% of the heritability in the three occupational status measures captured in our GWASs can be explained by genetic correlations with GWAS summary statistics of just these five traits. Between traits, associations are largely similar: The strongest effects are observable for cognitive performance. However, they are reduced by more than 30% once ADHD and particularly openness are added to the model. The influence of ADHD is then itself increased by the introduction of risk tolerance, whose effect on the SES factor is - in contrast to ADHD and neuroticism - positive. The latter is furthermore the only covariate with heterogeneous effects. While significantly negatively associated with SIOPS and almost significant for ISEI the neuroticism coefficient is markedly smaller and insignificant for CAMSIS.

| | (1) | (2) | (3) | (4) | (5) |
|---|---|---|---|---|---|
| Cognitive Performance | 0.695*** | 0.593*** | 0.437*** | 0.446*** | 0.440*** |
| | (0.021) | (0.029) | (0.071) | (0.071) | (0.069) |
| ADHD | | -0.292*** | -0.360*** | -0.412*** | -0.392*** |
| | | (0.040) | (0.067) | (0.073) | (0.076) |
| Openness | | | 0.366** | 0.340** | 0.338** |
| | | | (0.129) | (0.128) | (0.127) |
| Risk Tolerance | | | | 0.149** | 0.143** |
| | | | | (0.055) | (0.055) |
| Neuroticism | | | | | -0.069 |
| | | | | | (0.043) |
| R2 | 0.483 | 0.558 | 0.67 | 0.688 | 0.693 |
| *Note:* | | | | | **p<0.05; **p<0.01; ***p<0.001* |

*Supplementary Table 1. Results of multivariate genetic regression models - CAMSIS and potential mediators*

| | (1) | (2) | (3) | (4) | (5) |
|---|---|---|---|---|---|
| Cognitive Performance | 0.680*** | 0.590*** | 0.449*** | 0.459*** | 0.450*** |
| | (0.022) | (0.029) | (0.069) | (0.067) | (0.065) |
| ADHD | | -0.258*** | -0.319*** | -0.383*** | -0.348*** |
| | | (0.046) | (0.067) | (0.073) | (0.075) |
| Openness | | | 0.331** | 0.299* | 0.296* |
| | | | (0.124) | (0.122) | (0.119) |

| | | | | | |
|---|---|---|---|---|---|
| Risk Tolerance | | | | 0.184*** | 0.173*** |
| | | | | (0.052) | (0.051) |
| Neuroticism | | | | | -0.124** |
| | | | | | (0.044) |
| R2 | 0.462 | 0.521 | 0.613 | 0.641 | 0.655 |
| *Note:* | | | | | **p<0.05; **p<0.01; ***p<0.001* |

*Supplementary Table 2. Results of multivariate genetic regression models - SIOPS and potential mediators*

| | (1) | (2) | (3) | (4) | (5) |
|---|---|---|---|---|---|
| Cognitive Performance | 0.712*** | 0.616*** | 0.464*** | 0.473*** | 0.464*** |
| | (0.041) | (0.047) | (0.091) | (0.092) | (0.091) |
| ADHD | | -0.275*** | -0.341*** | -0.393*** | -0.364*** |
| | | (0.058) | (0.086) | (0.091) | (0.095) |
| Openness | | | 0.357* | 0.331* | 0.328 |
| | | | (0.167) | (0.169) | (0.168) |
| Risk Tolerance | | | | 0.149* | 0.140* |
| | | | | (0.062) | (0.062) |
| Neuroticism | | | | | -0.103 |
| | | | | | (0.055) |
| R2 | 0.507 | 0.573 | 0.68 | 0.699 | 0.708 |
| *Note:* | | | | | **p<0.05; **p<0.01; ***p<0.001* |

*Supplementary Table 3. Results of multivariate genetic regression models - ISEI and potential mediators*

### 10. Direct and indirect effects

In practice, polygenic scores might capture direct as well as (potentially noncausal) indirect genetic effects, the latter attenuating the genomic signal once the analysis is restricted to families,[46,47] reducing the utility for many practical purposes (i.e., Raben et al. 2021).[48] In particular, traits related to socioeconomic status tend to show a reduction of effect sizes in designs that allow to distinguish between direct and indirect effects.[49,50] To gauge the magnitude of these indirect effects, three designs were used.

#### 10.1 Parental Control design

Using the genotyped subsample of the NCDS, we extended the out-of -sample prediction outlined in 8.2 by restricting our data to all respondents for whom information of the paternal occupation at age 11 was available, again pooled all waves beginning at age 33 (N = 2,988, 2,972, 2,897, 2,746, 2,369 for CAMSIS and 3,019, 2,959, ,2890, 2,742, 2,363 at age 33, 42, 46, 50, 55) and fitted a linear model with and without controlling for parental SES (operationalized as paternal occupational status at age 11 - using highest parental education lead to similar results) after controlling for age, sex and the first 10 principal components. We then computed the ratio of the PGS coefficient from both models. Results are shown in Supplementary Table 4 below.

| Measure | N | Ratio (PRSice-2) | Ratio (SBayesR) | Ratio (SBayesR+MTAG) |
|---|---|---|---|---|
| CAMSIS | 18,086 | 0.747 | 0.747 | 0.793 |
| ISEI | 18,093 | 0.738 | 0.738 | 0.78 |
| SIOPS | 18,093 | 0.752 | 0.752 | 0.796 |

*Supplementary Table 4. Effect size reduction when controlling for parental SES*

#### 10.2 Adoption design

We conducted another GWAS analysis for occupational status, this time omitting the 3,398 (SIOPS, ISEI) to 3,414 (CAMSIS) participants of European descent from the UK Biobank who reported being adopted and had available occupational data. We used MTAG again (SIOPS, ISEI, CAMSIS, income, and education, excluding the adoptee sample) to enhance our discovery and SBayesR to optimize the PGS's predictive capacity. In comparison to our best-performing PGS, effect sizes reduced 23.3%, 22.6%, and 27.3% respectively ($R^2$= 0.043, 0.031, 0.027 for CAMSIS, ISEI, SIOPS).

#### 10.3 Sibling design

In order to identify direct genetic effects, we use the sibship sample from UK Biobank and compute fixed effects, subtracting family means from dependent and independent variables. Figure 5 in the main text displays the attenuation of the within-family signal measured by the ratio of the standardized beta coefficient of the respective best performing polygenic score from 8.2 in a model with family fixed effects to a baseline model without fixed effects. In both cases models control for age, gender and 10 principal components on 24,579 individuals for CAMSIS, 24,472 for ISEI and SIOPS, 36,265 for education and 31,851 for income. Confidence intervals were obtained using bootstrap.

#### 10.4 The sibling design and assortative mating

A possible explanation for the discrepancy between within-family estimates of SNP effects and GWAS estimates derived from unrelated individuals' samples could be the phenomenon of assortative mating. When the phenotypes of parents exhibit a correlation, it can result in the creation of long-range linkage disequilibrium among SNPs, even extending across different chromosomes. The subsequent increase in genetic variation in the population is not mirrored within family, leading to a reduction of effects between siblings.

Lee, Wedow, Okbay et al. (2018)[3] derive an approximation of the ratio of effect sizes for a causal SNP *j* in the population, $\beta_{BF,j}$, and within a sibship, $\beta_{WF,j}$ that can be expected as a result of a phenotypical spousal correlation *r*, when the number of loci *M* goes to infinity:

$$\lim_{M\to\infty} \frac{\beta_{WF,j}}{\beta_{BF,j}} = 1 - r\frac{(h^2-h^4r)}{1-h^4r} ,$$

where $h^2$ denotes the narrow sense heritability under assortative mating. Assuming a PGS to be a noisy measure of effect size weighted alleles of independent causal SNPs (or their LD-based proxies), this formula should provide an approximation to the expected PGS-effect reduction under the assumption of spousal phenotypic assortment, that can be estimated by plugging in plausible values for $h^2$ and *r* from the literature. The former can be found in the recent behavior genetics literature: Hootegem et al. (2023)[51] estimate the narrow sense heritability of SIOPS in Norway to be 0.38, Erola et al. (2022)[52] $h^2$ for ISEI in Finland as 0.42 and Marks (2017)[53] in Australia as 0.37.

Turning to *r*, Clark and Cummins (2022)[54] recently estimated the degree of assortative mating on occupational status in England from 1754-2021. Using a new database of 1.7 million marriage records they found it to be remarkably high: Correcting for measurement error, the groom-bride correlation equalled $r = 0.8$ with little to no variation over time.

Assuming $h^2 = 0.4$ and $r = 0.8$, $\lim_{M \to \infty} \frac{\beta_{WF,j}}{\beta_{BF,j}} = 0.75$. Then, assortative mating would explain a 25% effect reduction.

However, assortative mating can lead to downward bias in heritability estimates from twin models (Wolfram and Morris, 2023)[55] and notably only Erola et al. (2022)[52] correct for this, though only by using the phenotypic spousal correlation on educational attainment, so that even higher expected effect reductions (0.7 if $h^2 = 0.5$) might be plausible.

### 11. Polygenic score prediction over the life course

For SIOPS, ISEI and CAMSIS, the incremental R-square was calculated by looking at the increase in variance explained when adding the corresponding PGS to a regression including sex and 10 principal components on either of the three occupational status metrics for all five available measurements (Supplementary Figure 77). We used the genotyped subsample of the National Child Development Study (NCDS) for these analyses and phenotypic data at ages 33, 42, 46, 50 and 55. As in the pooled case, prediction for all time points highest for CAMSIS, with comparable performance for ISEI and SIOPS. Again, SBayesR outperforms PRSice2 and is furthermore improved by the combination with an MTAG predictor (using the same construction as specified in the pooled case). Highest incremental $R^2$-values are achieved at age 33 (up to 0.099, SE = 0.0076 for CAMSIS) and 55 (up to 0.93, SE = 0.0081 for CAMSIS), with a noticeable dip at age 42 (0.45, SE = 0.0056 for CAMSIS).

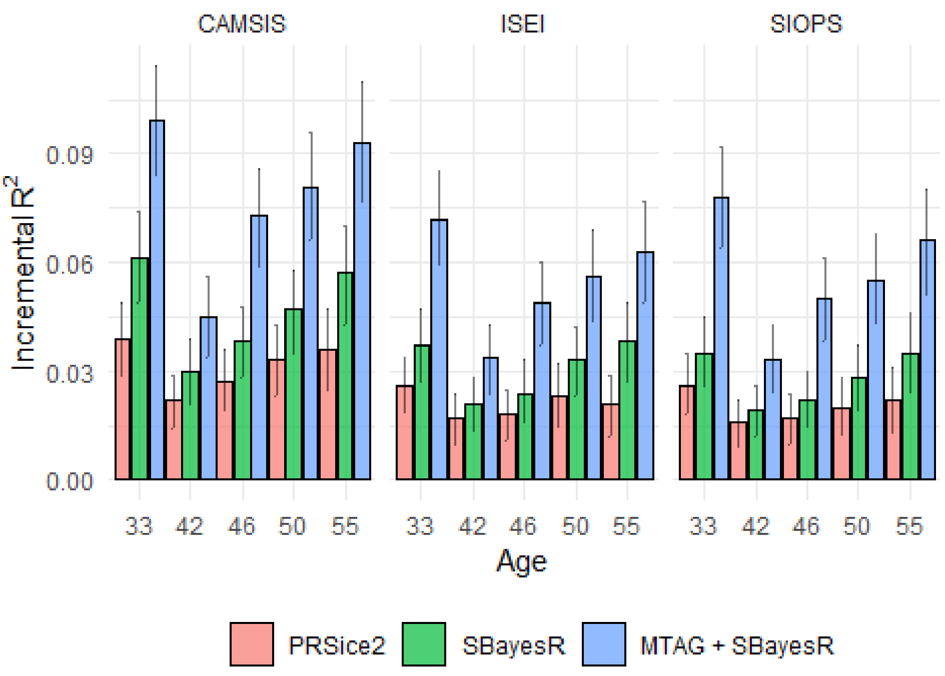


*Supplementary Figure 7. Incremental R-square of polygenic score predictions of occupational status over the life course, NCDS*

#### 12. Polygenic scores and the intergenerational transmission of occupational status

In addition to the various measures of occupational status over the life course, we also have information on the paternal occupation at age 11 in the NCDS data. The phenotypical correlation between paternal and offspring occupational status at the various ages for all three measures is substantial (r=0.3). However, while often interpreted as a purely social measure of the intergenerational transmission of occupational status, it might be confounded by shared genetic potential between father and child.

Using an approach first proposed by Tucker-Drob (2017),[37] we investigate the following scenarios. Specifically, what share of the intergenerational correlation for each of the three metrics at the beginning, middle and end of the career is confounded by the corresponding polygenic score if we assume that it only explains the amount of variance it does (1) or (2) it explains the full SNP-heritability? In the first case, a small but significant amount of confounding is observed for all metrics at all ages. In the second scenario 24-64% of the intergenerational correlation is confounded by genetic factors. Figure 8 in the main text displays the full results of these analyses.

#### 13. Polygenic score mediation by occupational aspirations and psychological traits

While we already showed that causally upstream psychological traits from the cognitive and noncognitive domain explain more than 70% of the SNP-heritability in occupational status, the NCDS data allows for the study of another potential pathway from genes to occupational status in the form of occupational aspirations. Respondents were asked at age 11 about the type of job they would like to do in the future. We coded these jobs to SOC2000, constructed ISEI, SIOPS and CAMSIS scores and ran mediation models from PGS to phenotypic occupational status, controlling for mediation by cognitive ability, internalizing behaviour, scholastic motivation, and externalizing behaviour in addition to aspirations. As with the genetic regression models, the largest share of the association is explained by cognitive ability. However, for all measures and ages, a significant share of the PGS (6.5-11%, Figure 7 in the main text) is mediated by aspirations.

To control for bias from indirect genetic effects in population estimates, we replicate these analyses controlling for parental socioeconomic status, determined by paternal occupational status at age 11. We find no substantial differences in our results (Supplementary Figure 8). Whilst the total effect's size decreases, as documented in Supplementary Information 10, the relative contributions of mediators largely persist, with only minor reductions in the mediating effect of cognitive ability accompanied by slight increases in the share of unexplained mediation.

### 14. Polygenic scores and occupational status trajectories throughout the life course

As occupational status and prestige measures are available at five consecutive time points in NCDS, we are also able to investigate to what extent PGS differences affect the trajectory of occupational status (N = 2,883 for CAMSIS); N = 2,894 for SIOPS/ISEI). For this, we ran growth curve models where allowing both the intercept and slope of CAMSIS, ISEI and SIOPS to vary over time as a function of the corresponding PGS. To allow more flexibility, quadratic terms for the slope were also added. We then iteratively removed effects of the PGS on squared slope, slope and intercept and compare model fit.

In all three cases, the baseline unconstrained model achieves the best fit, measured by AIC. In Supplementary Tables 5 and 6, we therefore report the coefficients for the unconstrained model. Effects of the PGS on occupational status trajectories are (highly) significant for all three status measures.

We also visualize the predicted status trajectories for all three metrics (Figure 6 in the main text). All show a comparable pattern: At the lower end of the PGS-distribution, status at the beginning of the career (age 33) is comparably low but increases rather steeply up for roughly a decade before then either stabilizing or even decreasing again. At the upper end of the distribution, however, a much higher starting prestige is followed by compounding status gains which show no sign of stopping up to age 55.

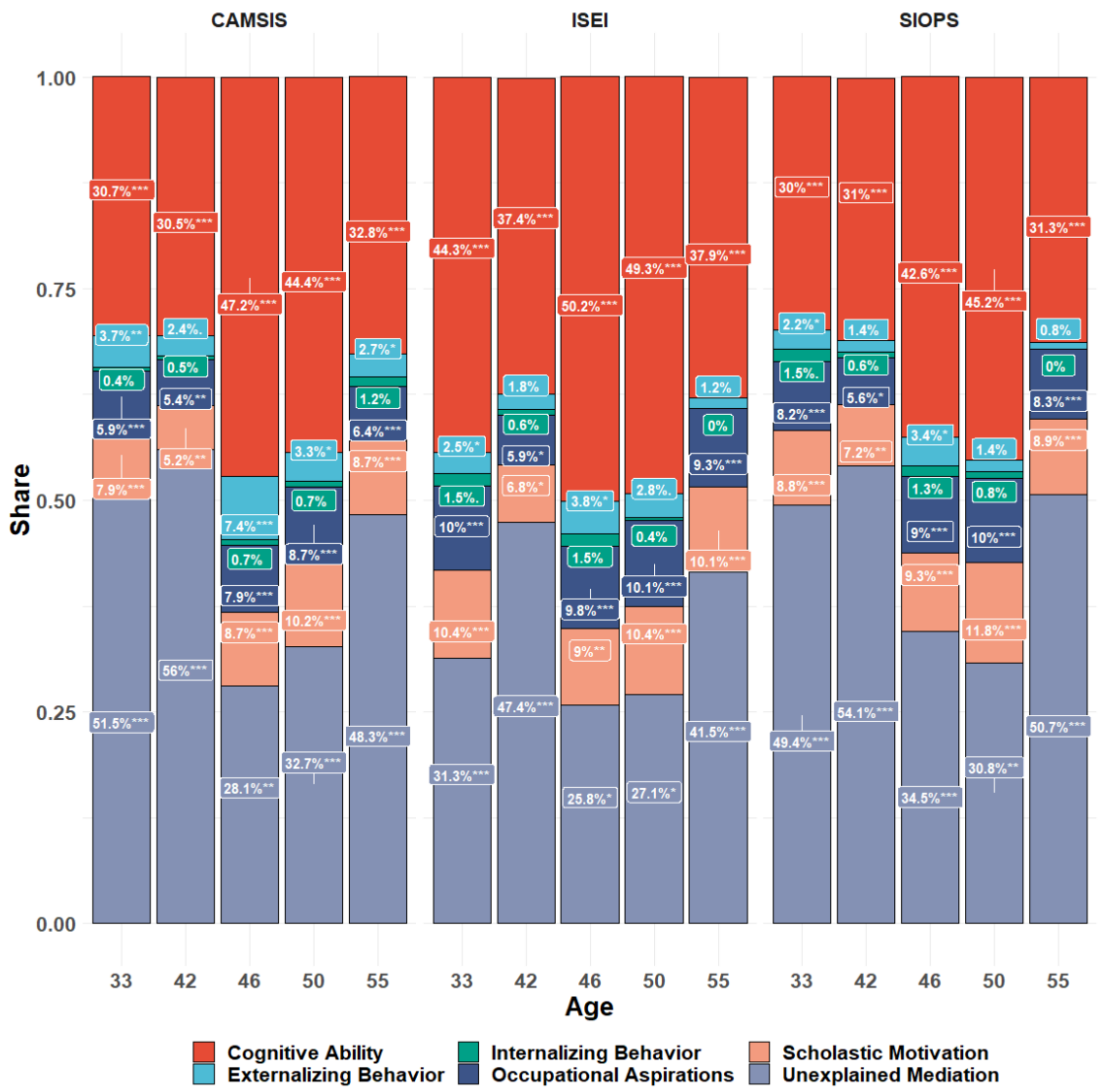


*Supplementary Figure 8. Mediation results of polygenic prediction of occupational status (controlling for parental SES), NCDS*

In addition, the robustness of this association is examined by accounting for parental socioeconomic status, quantified through the father's occupational status at the age of 11. The influence of all three status indicators is reducing controlling for such indirect effects. With respect to CAMSIS, the relationship between the PGS and the intercept, linear slope, and quadratic slope maintains its significance. PGS effect reduction and the association between the growth curve parameters and SES is comparable for SIOPS and ISEI in magnitude, however, neither PGS nor SES show a significant effect on the slope, a result best explained by the collinearity between PGS and SES combined with the lower predictive power of the SIOPS and ISEI polygenic scores.

| | CAMSIS | ISEI | SIOPS |
|---|---|---|---|
| Intercept at PGS = 0 | 52.8716***<br>(0.2382) | 44.8166***<br>(0.2914) | 42.5394***<br>(0.2406) |
| Slope at PGS = 0 | 0.2229***<br>(0.0276) | 0.192***<br>(0.0361) | 0.1556***<br>(0.0304) |
| Squared Slope at PGS = 0 | -0.0044***<br>(0.0012) | -0.0045**<br>(0.0015) | -0.0033**<br>(0.0012) |
| Effect of PGS on Intercept | 3.3094***<br>(0.2411) | 3.0265***<br>(0.2924) | 2.5205***<br>(0.2434) |
| Effect of PGS on Slope | -0.1491***<br>(0.0279) | -0.0724*<br>(0.0363) | -0.0657*<br>(0.0307) |
| Effect of PGS on Squared Slope | 0.007***<br>(0.0012) | 0.004**<br>(0.0015) | 0.0035**<br>(0.0013) |

*Supplementary Table 5. Parameters of Best Fitting Growth Curve Models, NCDS*

| | CAMSIS | ISEI | SIOPS |
|---|---|---|---|
| Intercept at PGS & SES = 0 | 52.6552*** (0.2713) | 44.5571*** (0.3357) | 42.5734*** (0.2815) |
| Slope at PGS & SES = 0 | 0.2079*** (0.0324) | 0.1923*** (0.0428) | 0.1399*** (0.036) |
| Squared Slope at PGS & SES = 0 | -0.004** (0.0014) | -0.0053** (0.0018) | -0.0035* (0.0015) |
| Effect of PGS on Intercept | 2.3402*** (0.2756) | 2.052*** (0.3373) | 2.0005*** (0.284) |
| Effect of SES on Intercept | 3.3084*** (0.2654) | 4.0552*** (0.3195) | 2.6632*** (0.2777) |
| Effect of PGS on Slope | -0.1077** (0.033) | -0.0442 (0.043) | -0.0358 (0.0363) |
| Effect of SES on Slope | -0.0713* (0.0317) | -0.035 (0.0407) | -0.0149 (0.0355) |
| Effect of PGS on Squared Slope | 0.0055*** (0.0014) | 0.0034 (0.0018) | 0.0027 (0.0015) |
| Effect of SES on Squared Slope | 0.0033* (0.0013) | 9e-04 (0.0017) | 7e-04 (0.0014) |

*Supplementary Table 6. Parameters of Best Fitting Growth Curve Models (controlling for parental SES), NCDS*

### 15. Polygenic score associations with health outcomes

In addition to the associations between the polygenic signals and occupational status, we also study its association with mental and general health outcomes. We show that the occupational status PGS has weak but robust links to health over the life course.

#### 15.1 General health

Within the NCDS, we look at health measured at ages 23, 33, 42, 46, 50 and 55. Participants were asked to rate their general health on a scale from:

- one (excellent) to four (poor) (age 23 and 33)
- one (excellent) to five (very poor) (age 42),
- one (excellent) to five (poor) (age 46, 50 and 55)

For each time point, the outcome is treated as metric and standardized to have a mean of zero and a standard deviation of 1. We then regress it on the CAMSIS, ISEI and SIOPS PGSs, respectively, while controlling for sex and ten principal components to correct for population stratification. We find weak but highly significant positive associations of the polygenic scores in all analytic models. Results are presented in Supplementary Table 7.

| | CAMSIS | ISEI | SIOPS | N |
|---|---|---|---|---|
| General Health at Age 23 | -0.058*** (0.013) | -0.054*** (0.013) | -0.052*** (0.013) | 5,586 |
| General Health at Age 33 | -0.101*** (0.013) | -0.093*** (0.013) | -0.088*** (0.013) | 5,686 |
| General Health at Age 42 | -0.079*** (0.012) | -0.075*** (0.012) | -0.07*** (0.012) | 6,233 |
| General Health at Age 46 | -0.086*** (0.013) | -0.086*** (0.013) | -0.088*** (0.013) | 5,925 |
| General Health at Age 50 | -0.099*** (0.013) | -0.092*** (0.013) | -0.088*** (0.013) | 5,666 |
| General Health at Age 55 | -0.11*** (0.013) | -0.106*** (0.013) | -0.105*** (0.013) | 5,302 |

*Supplementary Table 7. Associations between occupational status PGS and general health at various ages, controlling for sex and first 10 PCs*

#### 15.1a Controlling for parental occupational status

Controlling for paternal occupational status at age 11 - and therefore partly for social stratification in the GWAS - does attenuate the associations; however, significant associations remain. Results are presented in Supplementary Table 8.

| | CAMSIS | | ISEI | | SIOPS | | N |
|---|---|---|---|---|---|---|---|
| | Estimate | Ratio | Estimate | Ratio | Estimate | Ratio | |
| General Health at Age 23 | -0.046** (0.015) | 0.77 | -0.043** (0.015) | 0.82 | -0.045** (0.015) | 0.81 | 4,050 |
| General Health at Age 33 | -0.078*** (0.015) | 0.77 | -0.077*** (0.015) | 0.83 | -0.079*** (0.015) | 0.83 | 4,065 |
| General Health at Age 42 | -0.06*** (0.015) | 0.73 | -0.062*** (0.014) | 0.82 | -0.065*** (0.014) | 0.82 | 4,441 |
| General Health at Age 46 | -0.052*** (0.015) | 0.71 | -0.064*** (0.015) | 0.86 | -0.068*** (0.015) | 0.87 | 4,229 |
| General Health at Age 50 | -0.071*** (0.015) | 0.75 | -0.069*** (0.015) | 0.80 | -0.074*** (0.015) | 0.83 | 4,067 |
| General Health at Age 55 | -0.078*** (0.016) | 0.76 | -0.082*** (0.016) | 0.84 | -0.09*** (0.016) | 0.86 | 3,804 |

*Supplementary Table 8. Associations between occupational status PGS and general health at various ages, controlling for father's occupational status at age 11, sex and first 10 PCs. Ratio denotes the ratio of the standardized beta-coefficient of the PGS without controlling for father's occupational status and the standardized beta-coefficient of the PGS when controlling for father's occupational status in the same sample of individuals for which paternal occupational status information was available.*

#### 15.1b Confounding of phenotypical effect of occupational status

The association between phenotypical occupational status and general health is only weakly confounded by the polygenic signal. Even when we scale the polygenic score to the SNP-heritability using the method proposed in section 12, at maximum one third of the association is confounded by the polygenic score (Supplementary Figure 9).

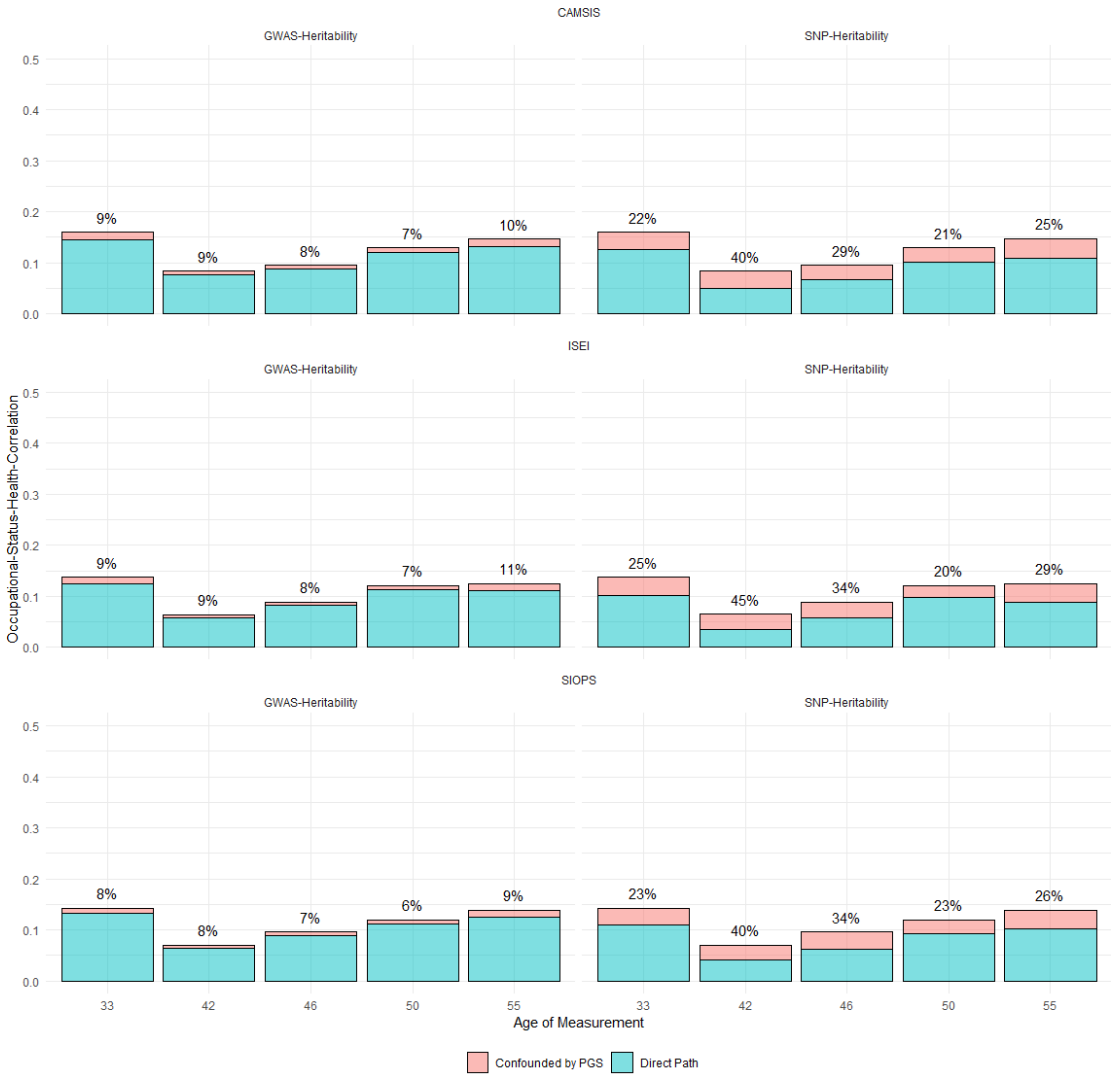


*Supplementary Figure 9. Genetic confounding of correlations from respondent's occupational status to general health*

#### 15.2 Mental health

Mental health in the NCDS is measured using the Malaise Inventory,[56] a commonly used self-completion scale for the assessment of psychiatric distress. Validation studies (partially conducted within the NCDS) show high reliability and validity of the instrument.[57]

The instrument is comprised of 24 questions. Each item has the response options yes or no:

1. *Do you often have backache?*
2. *Do you feel tired most of the time?*
3. *Do you often feel miserable or depressed?*
4. *Do you often have bad headaches?*
5. *Do you often get worried about things?*

6. *Do you usually have great difficulty in falling or staying asleep?*
7. *Do you usually wake unnecessarily early in the morning?*
8. *Do you wear yourself out worrying about your health?*
9. *Do you often get in a violent rage?*
10. *Do people often annoy and irritate you?*
11. *Have you at times had twitching of the face, head, or shoulders?*
12. *Do you often suddenly become scared for no good reason?*
13. *Are you scared to be alone when there are no friends near you?*
14. *Are you easily upset or irritated?*
15. *Are you frightened of going out alone or of meeting people?*
16. *Are you constantly keyed up and jittery?*
17. *Do you suffer from indigestion?*
18. *Do you suffer from an upset stomach?*
19. *Is your appetite poor?*
20. *Does every little thing get on your nerves and wear you out?*
21. *Does your heart often race like mad?*
22. *Do you often have bad pains in your eyes?*
23. *Are you troubled with rheumatism or fibrositis?*
24. *Have you ever had a nervous breakdown?*

The Malaise Inventory was part of the survey at ages 23, 33, 42 and 50. However, at age 50, only items 3, 5, 9, 12, 14, 16, 20 and 21 were asked. We therefore restrict ourselves to these eight items in our analyses. For all four time points, composite scores are created by the means of factor analysis and subsequently standardized. Loadings on a single factor were overall very high and similar for the different time points.

For each wave, we regress the three occupational status polygenic scores separately on the factor, while controlling for sex and the first ten principal components. Here, we find weak but significant positive associations between mental health and the three PGSs. Results are presented in Supplementary Table 9.

| | CAMSIS | ISEI | SIOPS | N |
|---|---|---|---|---|
| Mental Health at Age 23 | -0.08*** (0.012) | -0.074*** (0.012) | -0.073*** (0.012) | 5,545 |
| Mental Health at Age 33 | -0.054*** (0.012) | -0.05*** (0.012) | -0.049*** (0.012) | 5,698 |
| Mental Health at Age 42 | -0.036** (0.012) | -0.036** (0.012) | -0.038** (0.012) | 6,205 |
| Mental Health at Age 50 | -0.033* (0.014) | -0.042** (0.014) | -0.035* (0.014) | 5,057 |

*Supplementary Table 9. Associations between occupational status PGS and mental health at various ages, controlling for sex and first 10 PCs*

#### 15.2a Controlling for parental occupational status

Supplementary Table 10 presents the results of the models where we control for paternal occupational status. Some of the results are not significant but remain directionally similar which means that parental characteristics confound the association between occupational status polygenic score and mental health.

| | CAMSIS | | ISEI | | SIOPS | | N |
|---|---|---|---|---|---|---|---|
| | Estimate | Ratio | Estimate | Ratio | Estimate | Ratio | |
| Mental Health at Age 23 | -0.056*** (0.014) | 0.80 | -0.05*** (0.014) | 0.86 | -0.05*** (0.014) | 0.84 | 4,018 |
| Mental Health at Age 33 | -0.034* (0.015) | 0.70 | -0.027. (0.015) | 0.74 | -0.027. (0.015) | 0.72 | 4,072 |
| Mental Health at Age 42 | -0.029* (0.014) | 0.72 | -0.029* (0.014) | 0.77 | -0.032* (0.014) | 0.80 | 4,420 |
| Mental Health at Age 50 | -0.02 (0.016) | 0.64 | -0.032* (0.016) | 0.82 | -0.024 (0.016) | 0.75 | 3,646 |

*Supplementary Table 10. Associations between occupational status PGS and mental health at various ages, controlling for father's occupational status at age 11, sex and first 10 PCs. Ratio denotes the ratio of the standardized beta-coefficient of the PGS without controlling for father's occupational status and the standardized beta-coefficient of the PGS when controlling for father's occupational status in the same sample of individuals for which paternal occupational status information was available.*

#### 15.2b Confounding of phenotypical effect of occupational status and mental health

The association between phenotypical occupational status and general health is only weakly confounded by the polygenic signal. Once we scale the polygenic score to the SNP-heritability, the confounding becomes stronger. Nevertheless, the overall effects are small, as seen in Supplementary Figure 10.

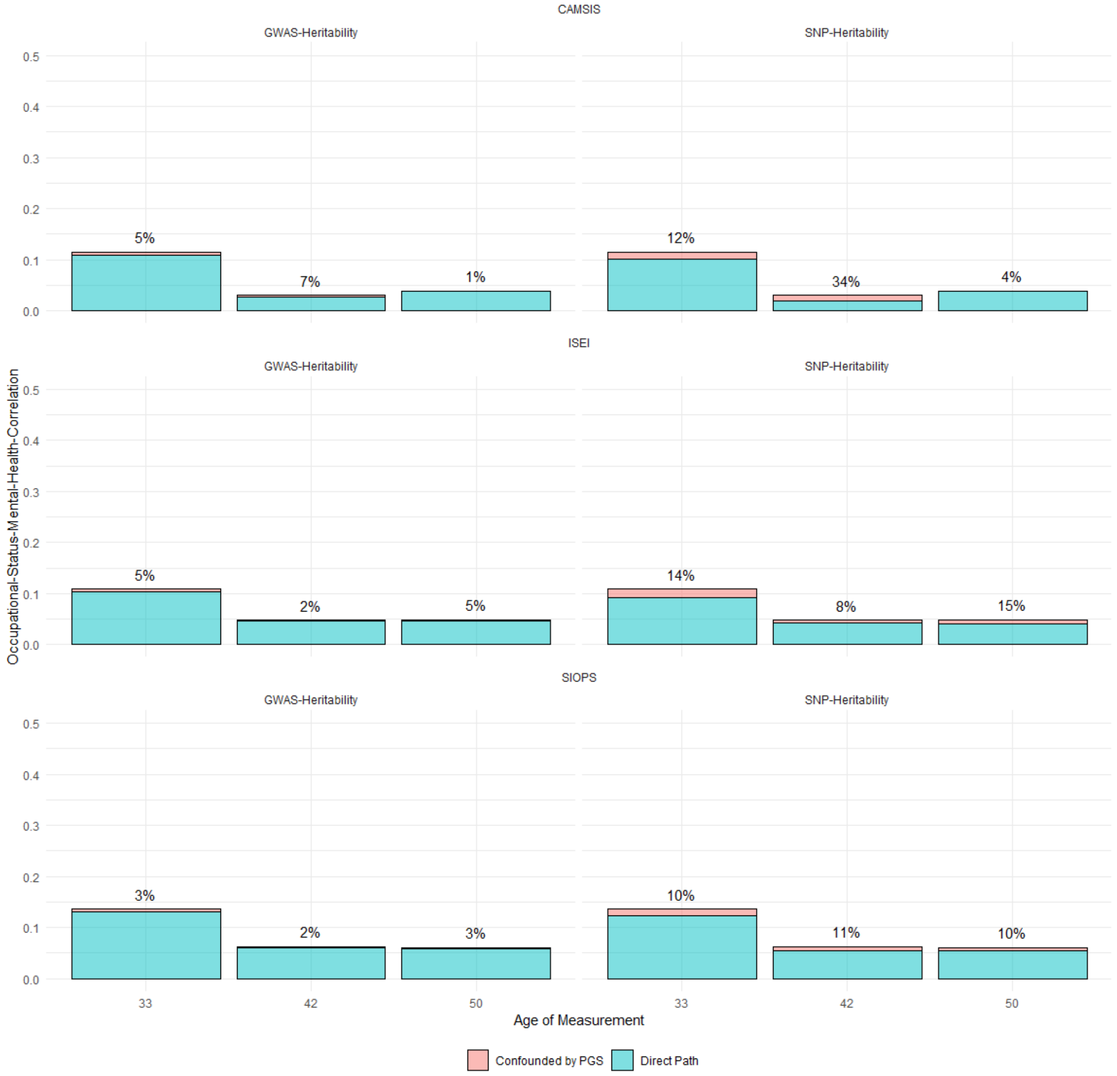


*Supplementary Figure 10. Genetic confounding of correlations from respondent's occupational status to mental health*
